# Caleosin 1 contributes to seed lipid droplet degradation by interaction with autophagy-related protein ATG8

**DOI:** 10.1101/2023.04.04.535563

**Authors:** Magdalena Miklaszewska, Krzysztof Zienkiewicz, Ewa Klugier-Borowska, Marcin Rygielski, Ivo Feussner, Agnieszka Zienkiewicz

## Abstract

Triacylglycerols (TAGs) accumulate in lipid droplets (LDs) of seed tissues to provide energy and carbon for seedling establishment. In the major route of LD degradation (lipolysis), TAGs are mobilized by lipases. However, LDs may be also degraded *via* lipophagy, a type of selective autophagy, which mediates LDs delivery to vacuoles or lysosomes. The exact mechanism of this process in plants still remains unresolved. Here, we provide evidence that during *Arabidopsis thaliana* seed germination, LDs are degraded by microlipophagy and that this process requires caleosin 1 (CLO1), a LD surface protein. We show co-localization of autophagy-related protein 8b (ATG8b) and LDs during seed germination and localization of lipidated ATG8 (ATG8-PE) to the LD fraction. We further demonstrate that CLO1, CLO2 and CLO3 interact with ATG8 proteins *via* their ATG8-interacting motifs (AIMs). Deletion of AIM localized directly before the proline knot disrupts CLO1 interaction with ATG8b, suggesting the essential role of this region in the interaction between the two proteins. Collectively, we provide new insights into the molecular mechanisms governing the interaction of LDs with the autophagy machinery in plant cells, contributing to understanding of the role of structural LD proteins in lipid mobilization.

## INTRODUCTION

Oilseed plants accumulate lipids mainly in the form of triacylglycerols (TAGs) in specialized organelles called lipid droplets (LDs) (Chapman and Ohlrogge, 2012; Xu and Shanklin, 2016; Miklaszewska et al., 2021). LDs consist of a neutral lipid core surrounded by a phospholipid monolayer harbouring a tissue-specific set of structural and associated proteins (Gidda et al., 2016; Huang, 2018; Fernández-Santos et al., 2020; Scholz et al., 2022). In oilseeds, the major structural proteins of LDs belong to three protein families: oleosins, caleosins, and steroleosins (Huang, 2018). Beside serving as a mere carbon and energy storage depot, LDs are actually dynamic structures involved in a plethora of developmental and physiological processes in plants, like seedling development, stress response, and pathogen resistance (Kelly et al., 2011; Blée et al., 2014; Fernández-Santos et al., 2020; Lu et al., 2020). LDs dynamics and function relies on the strictly controlled balance between their efficient synthesis and degradation in cells (Zienkiewicz and Zienkiewicz, 2020). During seed germination, the major route of LD degradation (lipolysis) involves lipases which hydrolyse TAGs into glycerol and free fatty acids (FFAs) (Graham, 2008). In *Arabidopsis thaliana,* SDP1 (SUGAR DEPENDENT1) has been shown to be responsible for breakdown of the majority of TAGs accumulated in the seeds (Eastmond, 2006; Kelly et al., 2011).

Recently, a prominent role of autophagy in lipid degradation has been described in eukaryotic cells (Seo et al., 2017; Fan et al., 2019). Autophagy is a conserved catabolic process, in which cellular components, including proteins, metabolites, and entire organelles, are digested within vacuoles (yeast, plants) or lysosomes (mammals) (Marshall and Vierstra, 2018; Allen and Baehrecke, 2020). A number of autophagy-related (ATG) proteins are directly involved in this process as core components of the cellular autophagic machinery. Among ATG proteins, ATG8 plays a prominent role in the autophagosome formation and serves as a docking site for cargos targeted for degradation by autophagy (Kellner et al., 2017; Bu et al., 2020). During this process, ATG8 proteins are lipidated by a phosphatidylethanolamine (PE) molecule *via* a ubiquitin-like conjugation system and anchored in the inner and outer membrane of the autophagosome (Li and Vierstra, 2012; Johansen and Lamark, 2020). The interaction between cargos targeted for autophagy-mediated degradation and the PE-conjugated ATG8 protein is mediated by the specific ATG8-interacting motifs present in autophagic receptors. Two types of ATG8 binding sites have been identified so far – AIMs/LIRs (ATG8-interacting motifs in plants and yeast / LC3-interacting regions in animals) and UIMs (ubiquitin interacting motifs) (Marshall et al., 2019). These motifs mediate the interaction of cargos with the AIM/LIR docking site (LDS) and the UIM docking site (UDS) of ATG8, respectively. AIM/LIR comprises of four amino acids: an aromatic amino acid (F/W/Y), followed by two random amino acids (X-X) and a branched chain amino acid (L/I/V) (Kellner et al., 2017; Johansen and Lamark, 2020). For UIMs, no consensus amino acid sequence has yet been identified due to a limited number of known ATG8-binding UIMs across species (Marshall et al., 2019).

Two general types of autophagy have been described – macroautophagy and microautophagy (Reggiori and Klionsky, 2013; Parzych and Klionsky, 2014; Wang et al., 2018). In the case of macroautophagy, cellular cargo (organelles or macromolecules) is encapsulated by double-membrane autophagosomes, which finally fuse with lysosomes/vacuoles for their cargo delivery and degradation. In microautophagy, on the other hand, cellular components are directly engulfed by lysosomes/vacuoles and then degraded (Parzych and Klionsky, 2014; Schuck, 2020). A type of selective autophagy directly involved in LD degradation has been termed lipophagy. In yeast, microlipophagy plays a predominant role in this process *via* ATG-dependent or -independent mechanisms (van Zutphen et al., 2014; Vevea et al., 2015; Seo et al., 2017). In mammalian cells, in addition to microlipophagy, a complete or “piecemeal” macrolipophagy is involved in LD turnover through *de novo* autophagosome formation followed by autophagic cargo degradation in the autolysosome (Ward et al., 2016; Khawar et al., 2021).

The most recent studies also suggest an important role of lipophagy in LD breakdown in algae and plants (Fan et al., 2019; Barros et al., 2020; Zienkiewicz et al., 2020). Microlipophagy-like degradation of LDs has been described for various microalgae like *Chlamydomonas reinhardtii*, *Auxenochlorella protothecoides* or *Nannochloropsis oceanica* (Zhao et al., 2014; Tsai et al., 2017; Zienkiewicz et al., 2020). In land plants, lipophagy seems to be involved in LD degradation during anther development and pollen germination (Kurusu et al., 2014; Zhao et al., 2020). A microlipophagy-resembling process has also been described during dark-induced starvation in Arabidopsis leaves (Fan et al., 2019). This process was impaired in Arabidopsis *atg2-1 sdp1-4* and *atg5-1 sdp1-4* suggesting a key role of core components of the macroautophagy machinery in LD degradation. Contribution of autophagy to LD degradation has also been suggested during seed germination in *A. thaliana* in a LD structural protein mutant – *caleosin1* (*Atclo1*) (Poxleitner et al., 2006). Loss of function of *At*CLO1 was associated with delayed LD mobilization when compared to wild type plants. Interestingly, unlike wild type plants, *Atclo1* mutants showed the absence of LD inside of vacuoles in cotyledon cells. Thus, it has been suggested that *At*CLO1 may be directly involved in microlipophagy during *A. thaliana* seed germination (Poxleitner et al., 2006).

Deciphering the molecular mechanisms behind LD degradation *via* lipophagy is still in its infancy, however, recent findings suggest the direct interaction of LD structural proteins with the autophagic machinery (Marshall et al., 2019; Zienkiewicz et al., 2020). Our previous studies showed that the LD surface protein (LDSP) from *N. oceanica* interacts with *No*ATG8 *via* its AIM (Zienkiewicz et al., 2020). Similarly, Marshall et al. (2019) showed that *At*OLE1 binds to the AIM/LIR docking site of *At*ATG8e. The fact that structural LD proteins are able to interact with ATG8 together with the observation that *clo1* mutants of *A. thaliana* are impaired in degradation of LDs inside the vacuoles (Poxleitner et al., 2006) prompted us to investigate whether caleosins mediate LD degradation by interaction with ATG8 and to decipher the mechanism of this process. To date, only a few of eight caleosins (CLO1-CLO8) identified in the Arabidopsis genome have been characterized (Næsted et al., 2000; Shen et al., 2014). All these proteins contain an N-terminal hydrophilic domain consisting of an EF-hand calcium binding motif, a central hydrophobic region with the proline knot that anchors the protein to the LD membrane and a C-terminal hydrophilic region with several phosphorylation- and heme-binding sites (Purkrtova et al., 2007; Chapman et al., 2012; Huang, 2018). The genes encoding CLO1 and CLO2 are highly expressed in developing seeds suggesting their role in seed germination or dormancy (Poxleitner et al., 2006; Liu et al., 2022). Consequently, we examined here in detail the phenotypic effects of a *CLO1* and *CLO2* mutation on LD degradation during *A. thaliana* seed germination and provide evidence on the interaction of these proteins with ATG8. Our study also demonstrates the essential role of CLO1 in LD degradation *via* microlipophagy during seed germination and reports novel data on the mechanistic basis of lipophagy in plants.

## RESULTS

### Characterization of *clo1* and *clo2* mutants

The *A. thaliana* genome contains eight caleosin genes (Shen et al., 2014). Two of them – *CLO1* (At4g26740) and *CLO2* (At5g55240) are expressed preferentially in developing seeds (Shen et al., 2014). To determine the function of Arabidopsis CLO1 and CLO2 in seeds, we characterized T-DNA insertion lines available for *CLO1* (*Atclo1-1*, Poxleitner et al., 2006) and *CLO2* (SALK_046559) in the Columbia 0 (Col-0) ecotype. PCR genotyping was performed to identify *clo1* and *clo2* homozygous mutant lines (Supplemental Figure S1A and B). Additionally, immunoblotting analyses showed that both *clo1* and *clo1 clo2* mutants failed to accumulate CLO1 protein in the mature seeds (Supplemental Figure S1C). To investigate the effects of the loss of function of CLO1 and CLO2 on seed morphology, we evaluated three different traits for seed yield in *clo1*, *clo2* and *clo1 clo2* mutants: seed dry weight, seed length and seed width (Supplemental Figure S2) and compared them to wild type plants (Col-0). The mature dry seeds obtained from the *clo1 clo2* double mutant were significantly longer and heavier, but not wider, when compared to both Col-0 and to the single caleosin mutants (Supplemental Figure S2A-2D). No significant changes in seed morphometry were observed between the single mutants and the control plants (Supplemental Figure S2A-2D).

### Disruption of CLO1- and CLO2-encoding genes affects acyl composition of mature and germinating seeds of Arabidopsis

To address the role of CLO1 and CLO2 in lipid metabolism, we first analysed acyl composition of total lipids from mature and germinating seeds of single and double mutants of *CLO1* and *CLO2* under diverse light conditions, and compared to the control plants (Figure 1, Supplemental Figure S3 and S4). In the mature seeds, the mutation in the CLO1-encoding gene was accompanied by a significant increase in 18:1 content and lower levels of 16:0 (Figure 1A). Moreover, all analysed *clo1* and *clo2* mutants showed significant reduction in 20:1. During the course of germination of all analysed mutants under long day conditions, a similar pattern of 18:1 and 20:1 content was observed, with exception to 96 h of germination, where no significant changes of the acyl content were found between *clo1*, *clo2* and *clo1 clo2* seedlings in comparison to Col-0 plants (Supplemental Figure S3). Similar changes in 18:1 and 20:1 levels were observed during *in vitro* germination under continuous dark (Supplemental Figure S4). As in Arabidopsis seeds 20:1 is found almost exclusively in TAGs, it has been commonly used as a marker of TAG degradation (Poxleitner et al., 2006; Kelly et al., 2011). Consequently, to determine if the *clo1* and *clo2* mutants are impaired in TAG breakdown, we analysed the changes in their 20:1 content during *in vitro* seed germination under long day (Figure 1B) and continuous dark (Figure 1C) conditions. In both cases, a progressive depletion of 20:1 was observed for all analysed plants, however, less obvious for seeds germinating under continuous dark. Under long day conditions, the content of 20:1 was significantly higher in plants with mutated *CLO1* (*clo1* and *clo1 clo2* plants), when compared to *clo2* and Col-0 (Figure 1B). Under continuous dark, however, the content of 20:1 was significantly higher in *clo1* and *clo1 clo2* seeds only after 72 and 96 h of germination (Figure 1C). Overall, the breakdown of 20:1 was slower during *in vitro* germination in the dark, independently from the *CLO1* and *CLO2* mutations. However, mutation in *CLO1* results in a slight delay of this process regardless of the light conditions.

**Figure 1.**
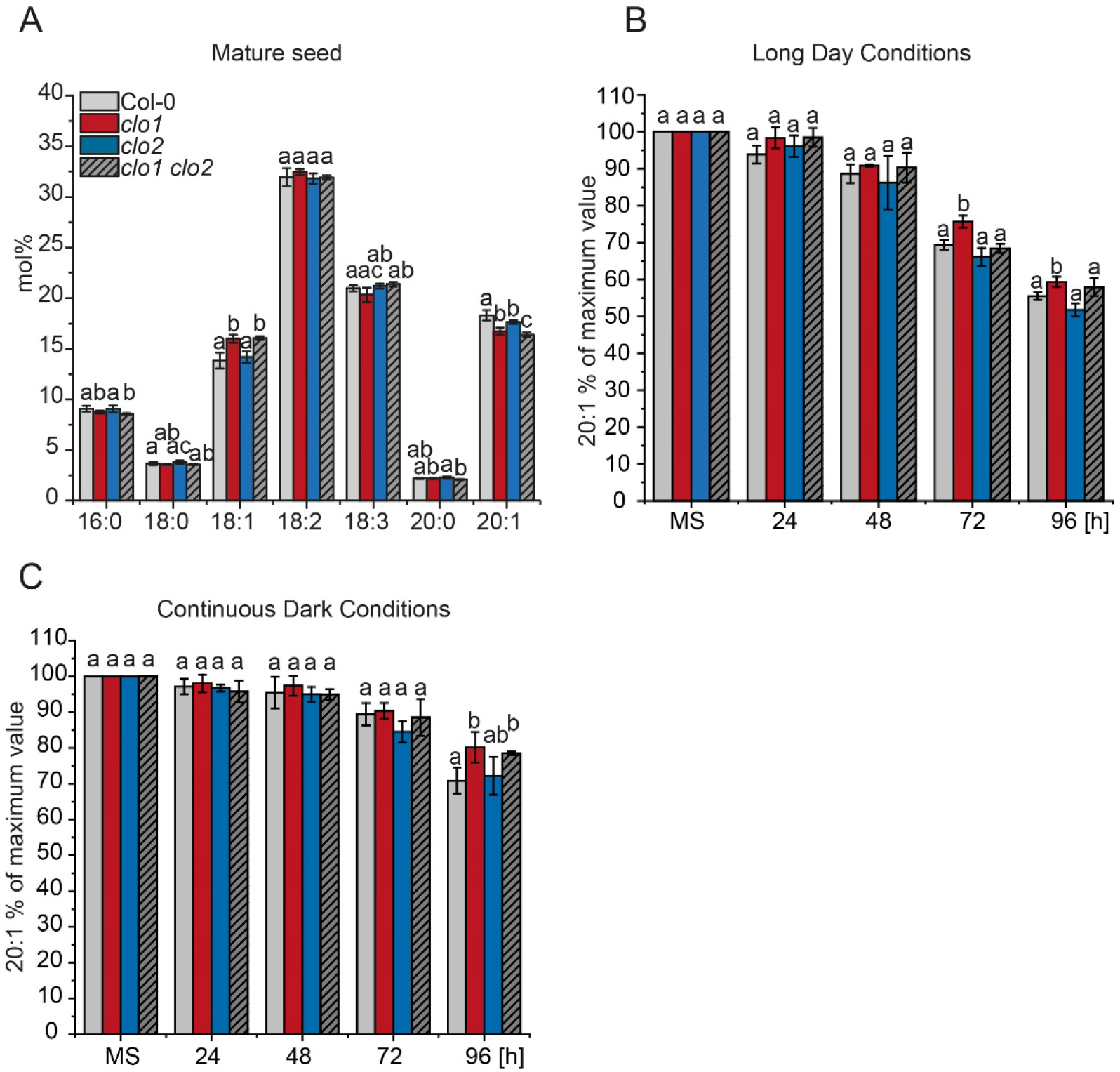
Lipid analyses of wild type plants and caleosin mutants during seed germination. (A) Changes in mature seed (MS) FAs composition between Col-0, *clo1*, *clo2* and *clo1 clo2*. (B-C) Changes in 20:1 content is presented as a percentage of the amount determined in the same number of mature seeds. The analysis was performed during germination course under long day (B) and continuous dark (C) conditions. Data are means ± SD from two independent experiment with six biological replicates (n = 6). The experiment was repeated three times with similar results using independent biological samples. Statistical analysis was performed by one-way ANOVA with Tukey‘s post hoc test. Different letters indicate significant differences with P < 0.05.

### Cellular organization of *clo1* mutant reveals disrupted LD breakdown

To confirm the impact of *CLO1* and *CLO2* mutations on TAG degradation during seed germination, we analysed cellular organization of LDs in cotyledons of *clo1*, *clo2* and *clo1 clo2* seedlings with reference to Col-0 by using confocal laser scanning microscopy (CLSM) and transmission electron microscopy (TEM) (Figures 2 and 3). At 48 h of *in vitro* germination under long day conditions, in both Col-0 and *clo2* cotyledons LDs were found to be randomly distributed throughout the cells between the forming chloroplasts (Figure 2A-D and I-L). In contrast, in the cells of *clo1* and *clo1 clo2* mutants, we found that LDs are usually present in the peripheral parts of the cells and are often accumulated together, forming clumps. As expected, under continuous dark, in all analysed lines no chloroplasts could be found at 48 h of germination. However, similar to long day conditions, evident changes in LD behaviour were found for single and double *clo1* mutants when compared to Col-0 and *clo2*. In the latter case, numerous LDs localized in both the periphery and centre of the cells, whereas in the vast majority of *clo1* and *clo1 clo2* cotyledon cells, LDs were mostly concentrated in the peripheral area of the cell around the central vacuole (Figure 2). To gain a more detailed insight into the cellular effects of *CLO1* and *CLO2* disruption in seeds germinating under long day conditions, we analysed the ultrastructure of *clo1*, *clo2* and *clo1 clo2* cotyledon cells, with reference to Col-0 (Figure 3). After 48 h of germination, in the cotyledon cells of Col-0 the most prominent feature was the presence of LDs at the border as well as inside the central vacuole (Figure 3A-D). Moreover, these LDs were usually surrounded by a thin layer of cytoplasmic material (Figure 3A-B, arrows). At the later steps of germination, in most of the cells, partially degraded LDs and residues of lipidic material were found in the central vacuole (Figure 3C). At the same time, the cytosolic pool of LDs was strongly depleted and composed mostly of small LDs (Figure 3C-D). Strikingly, at 48 h of *clo1* germination practically no LDs were observed in the central vacuole (Figure 3E-F). Instead, they were found as tightly packed structures in the cytosol, forming aggregates and pushing towards the tonoplast. In the case of *clo2* plants, there were no obvious differences in the ultrastructure compared with Col-0 as the LDs were present inside of the central vacuole and were also carrying cytosolic material on their surface (Figure 3G, arrow). Finally, the double mutation of *CLO1* and *CLO2*, similarly to *clo1* ultrastructural phenotype, was accompanied by the presence of numerous large LDs in the cytoplasm (Figure 3H). No LDs were observed inside the central vacuole of *clo1 clo2* cotyledons during the whole course of germination. These results show that the disruption of *CLO1*, but not of *CLO2*, is associated with the impaired LD entering to and breakdown in the central vacuole during Arabidopsis seed germination.

**Figure 2.**
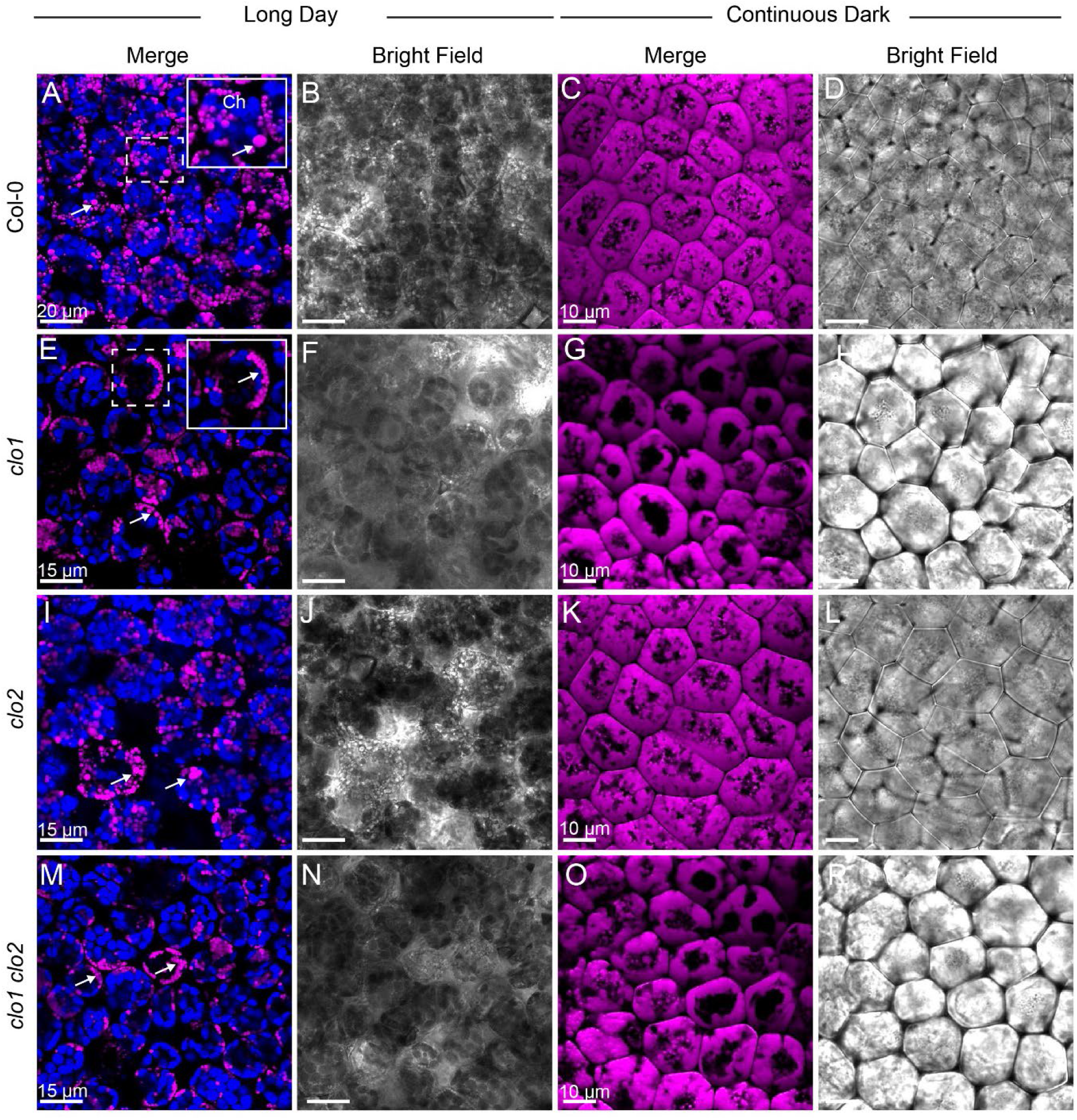
Microscopic analyses of LDs in germinating seeds of wild type plants and caleosin mutants. Representative CLSM images of cotyledon cells of Col-0 (A-D), *clo1* (E-H), *clo2* (I-L), (M-P) *clo1 clo2* after 48 h of germination under long day and continuous dark conditions showing LDs stained with BODIPY^TM^ 505/515 (arrows, magenta) and chlorophyll (blue).

**Figure 3.**
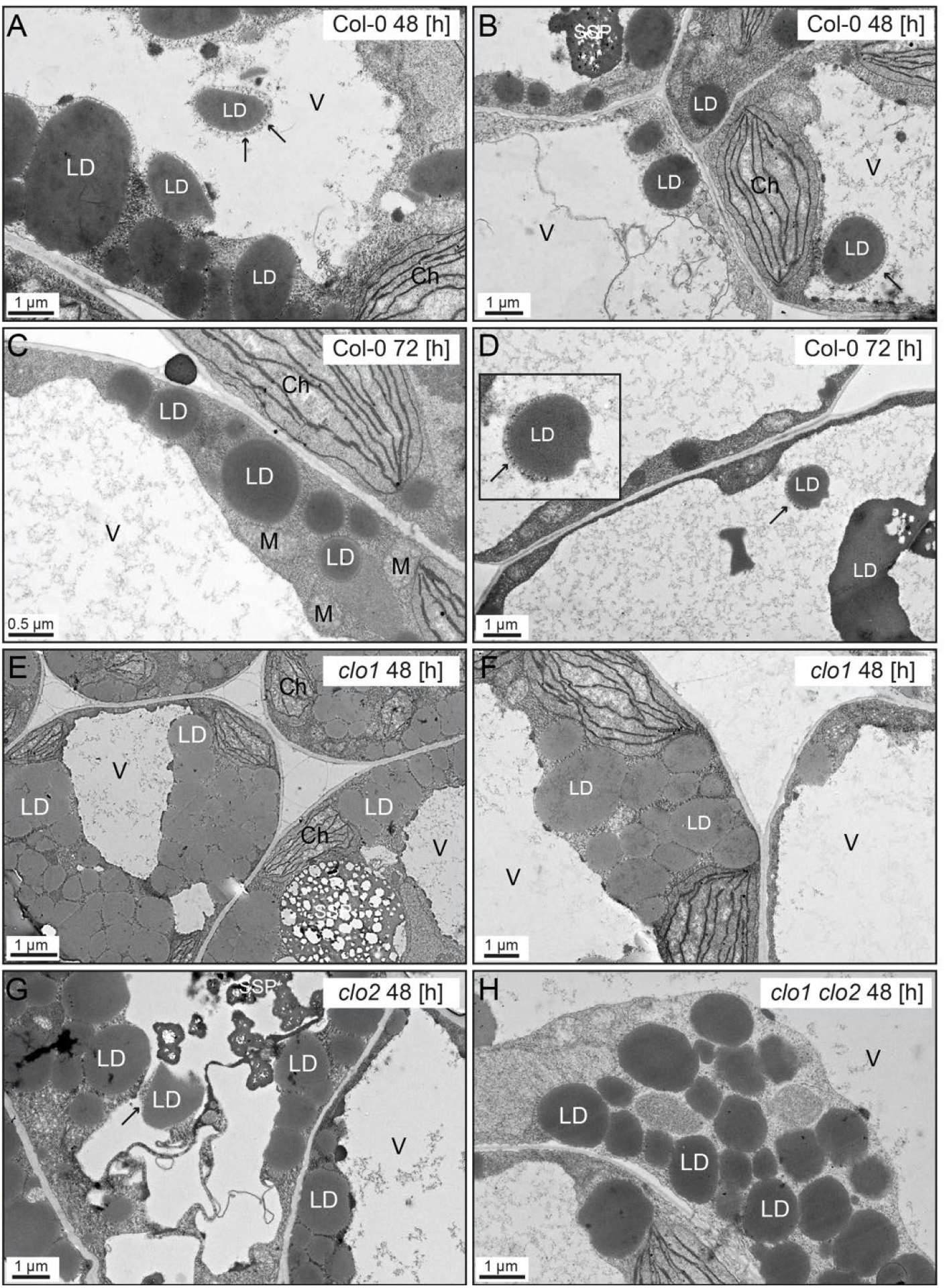
Ultrastructure of cotyledon cells during seed germination. Representative transmission electron microscopy (TEM) images of Arabidopsis cotyledons cells. (A-D), Ultrastructure of Col-0 cotyledons cells after 48 h (A-B) and 72 h (C-D) of *in vitro* germination. Lipid droplets (LDs) are present in the cytoplasm as well as inside the vacuole. The LDs visible in the area of the vacuole are surrounded by cytoplasmic material (arrows). (E-F), Ultrastructure of *clo1* cotyledons cells after 48 h of *in vitro* germination. LDs are localized only in the cytoplasm around the vacuole. (G), Ultrastructure of *clo2* cotyledons cells after 48 h of *in vitro* germination. LDs can be seen in the cytoplasm and inside the vacuole. (H), Ultrastructure of *clo1 clo2* cotyledons cells after 48 h of *in vitro* germination. LDs occupy only the cytoplasmic area. Ch, chloroplast; LD, lipid droplet; M, mitochondrion; SSP, seed storage protein; V, vacuole.

### In wild type Arabidopsis the vacuolar breakdown of LDs occurs in a process that resembles microlipophagy

To gain more information on the cellular mechanisms of LD degradation, we labelled the LDs with BODIPY^TM^505/5015 in transgenic *A. thaliana* plants overexpressing a tonoplast marker protein fused to cyan fluorescent protein (vac::CFP). Then, we analysed the behaviour of LDs during the course of seed germination by using CLSM (Figure 4). After 36 h (Figure 4 A-L) and 48 h (Figure 4 M-Z) of seed germination under long day conditions, we detected individual LDs inside the vacuoles. These LDs were often associated with invaginations (36 h) or directly surrounded (48 h) by the CFP-labelled tonoplast (Figure 4, arrows). Moreover, we observed the similar localization pattern for LDs during microscopic analysis of the plants expressing the CLO1-EYFP construct (Supplemental Figure S5). In cotyledon cells of germinating seeds, during early stages of seed germination (24 h), the CLO1-EYFP signal was observed in the form of numerous rings located in the cytoplasm (Supplemental Figure S5, arrowheads). However, the later steps of seed germination (48 h) were accompanied by a rich pool of these structures present in the area of the central vacuole (Supplemental Figure S5, arrows). Our observations strongly suggest that degradation of at least a certain number of LDs during seed germination is mediated by microlipophagy.

**Figure 4.**
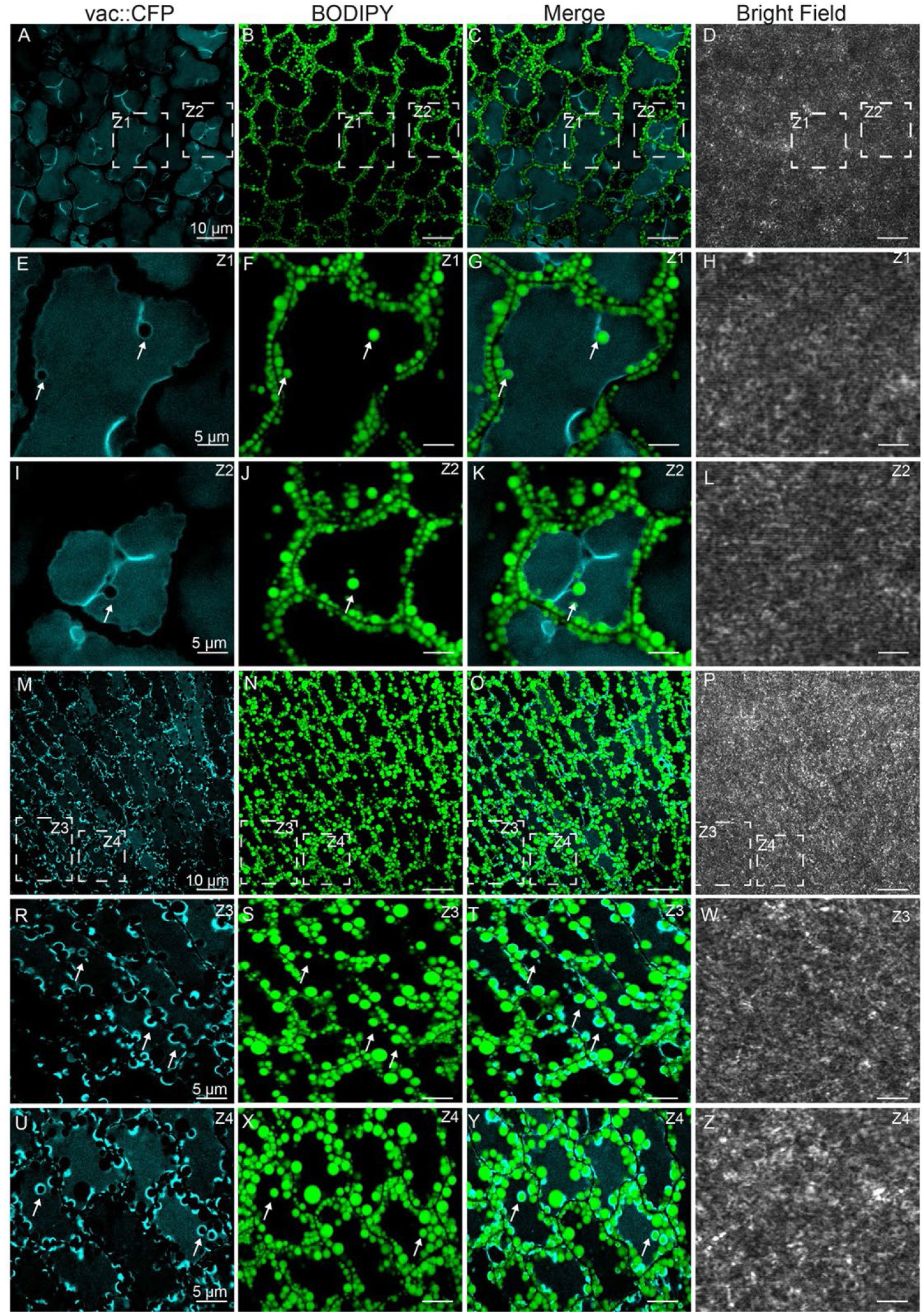
Microlipophagy of LDs during seed germination. Representative CLSM images of cotyledon cells of Col-0 after 36 h (A-L) and 48 h (M-Z) of seed germination under long day conditions. LDs were stained with BODIPY^TM^ 505/515 (green). Tonoplast marker protein fused to cyan fluorescent protein (CFP) is visible as blue. Arrows indicate the LDs within invaginations or completely surrounded by CFP-labelled tonoplast. The areas marked with dashed lines (Z) represent zoomed views of the corresponding image.

### ATG8 protein is associated with LDs during seed germination

To examine whether autophagy plays an essential role in LD degradation, we analysed a possible interaction between ATG8b and LDs (Figure 5). We confirmed *in situ* the co-localization of ATG8b with LDs in the cotyledon cells (Figure 5). By using plants stably expressing ATG8b-EYFP (Supplemental Figure S6), we demonstrated that in cotyledons ATG8b co-localizes with LDs during seed germination under both long day (Figure 5 A-P, arrows) and continuous dark conditions (Figure 5 R-W, arrows). This co-localization was observed in the mesophyll cells as well as in the stomata (Figure 5 M-P). By immunoblotting, we determined the presence and abundance of CLO1 and ATG8 proteins during Col-0 seeds germination (Figure 6A and B, respectively). An anti-CLO1 antibody detected a single protein band of ∼26 kDa at all analysed time points (Figure 6A). The highest levels of CLO1 were observed in total protein extracts from the mature seeds. From imbibition onwards, CLO1 levels progressively decreased, to its almost complete disappearance after 72 h of *in vitro* germination (Figure 6A). This protein pattern of CLO1 nicely correlated with the progress of LD degradation during seed germination and seedling development. By using an anti-ATG8 antibody, we were able to detect two pools of ATG8 protein in Col-0 germinating seeds: free ATG8 and ATG8-PE (Figure 6B). The two corresponding protein bands were the most prominent in the mature and imbibed seeds as well as after 24 h of germination. From 48 h onwards, a gradual depletion in both pools of ATG8 protein was observed, with the lowest levels at 96 h of germination (Figure 6B). To examine whether CLO1 and ATG8 are indeed associated with LDs during seed germination, we employed the two above-mentioned antibodies to determine the presence of both proteins in the protein extracts isolated from a LD fraction obtained after 24 h of germination from Col-0, *clo1*, *clo2* and *clo1 clo2* seeds (Figure 6C-E). As expected, specific protein bands corresponding to CLO1 were detected in LD fractions isolated from Col-0 and *clo2* seeds but not in those extracted from *clo1* and *clo1 clo2* seeds (Figure 6C). In the case of ATG8 and ATG8-PE, the most prominent band observed in the LD fractions extracted from all analysed plant lines corresponded to ATG8-PE, whereas ATG8 was detected at much lower level (Figure 6D). We further confirmed the distinct levels of ATG8 and ATG8-PE in the LD fraction by comparing their pattern with the total protein extracted from Col-0 rosette leaves (Figure 6E). Immunoblotting with the anti-ATG8 antibody revealed the presence of both ATG8 pools in the total protein fraction, but mainly ATG8-PE was detected in the LD fraction. Thus, the findings from these experiments suggested that mostly ATG8-PE is associated with LDs and that this association is unaffected by *CLO1* and *CLO2* mutations.

**Figure 5.**
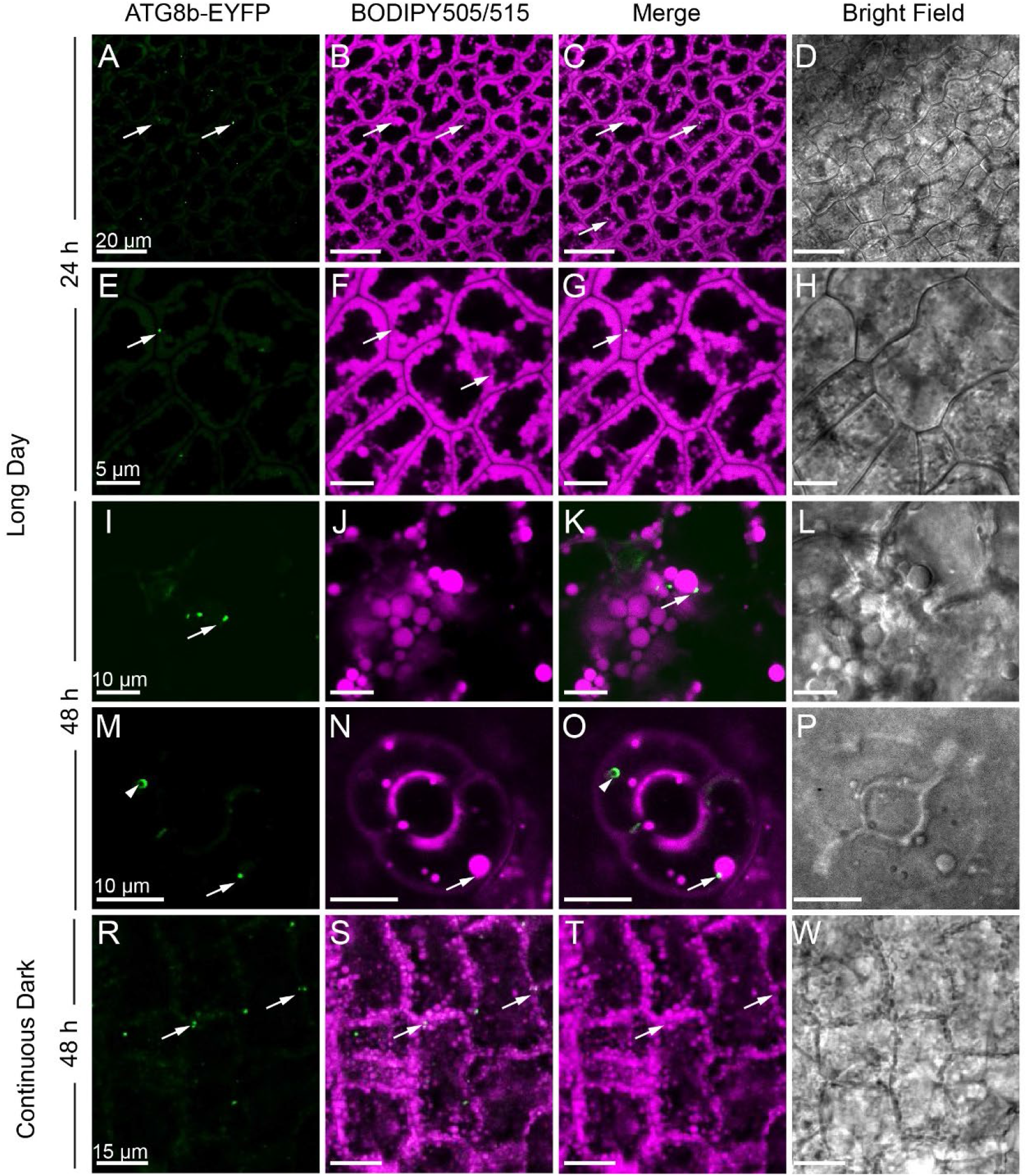
Co-localization of ATG8b and LDs. Representative CLSM images of cotyledon cells of ATG8b-EYFP transgenic line (green, arrows) stained with BODIPY 505/515 (magenta) and analysed after 24h (A-H) and 48 h (I-P) of seed germination under long day conditions as well as after 48 h (R-W) under continuous dark. Arrows indicate ATG8b-EYFP co-localizing with LDs.

**Figure 6.**
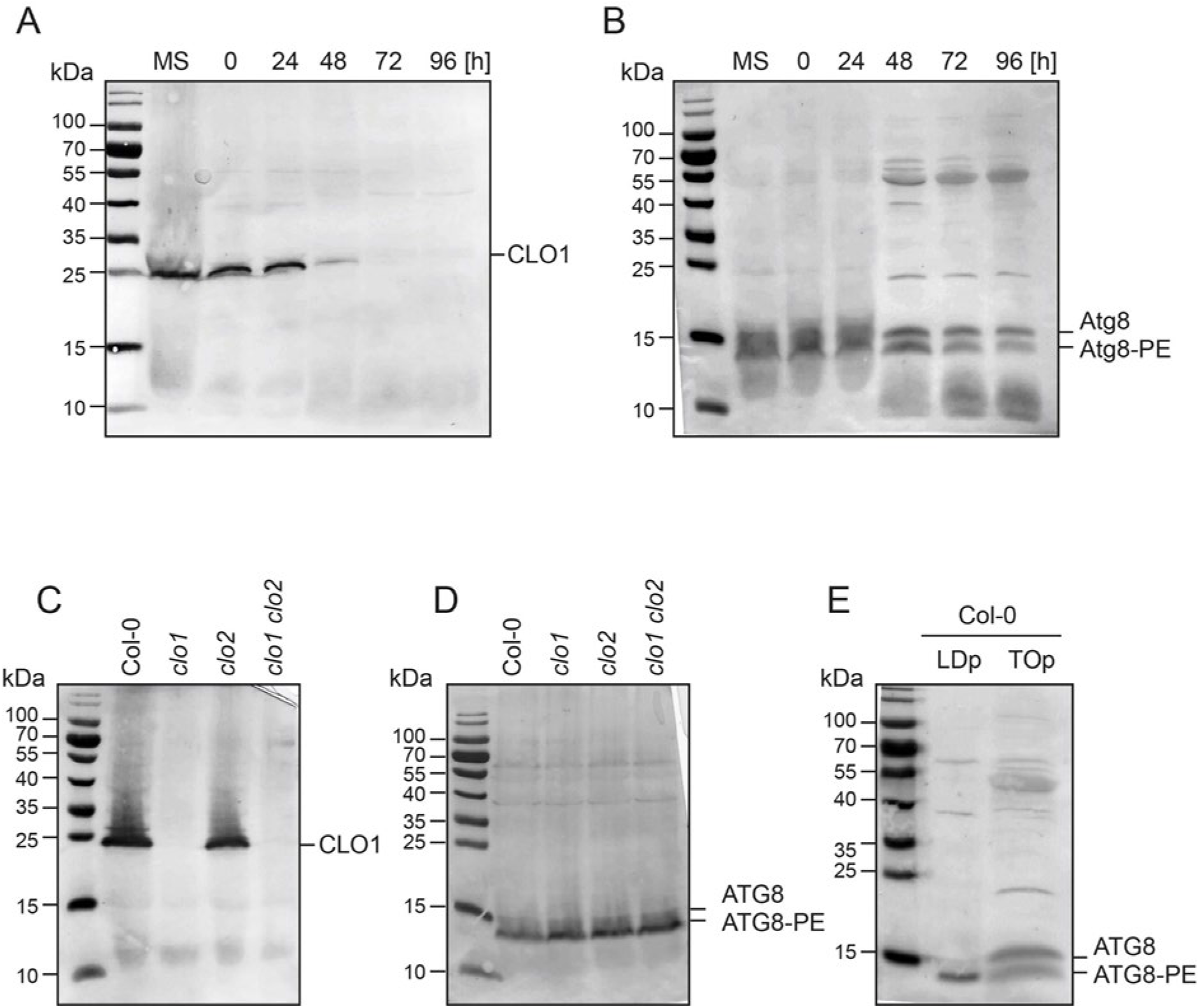
Caleosin and ATG8 abundance in wild type plants and caleosin mutants. (A-B) Immunoblot analysis of CLO1 and ATG8b levels during seed *in vitro* germination, respectively. Representative immunoblotting for CLO1 (C) and ATG8 (D) detected in LDs protein fraction extracted at 24 h of seed *in vitro* germination. (E) Detection of ATG8 and ATG8-PE pool in the total protein fraction extracted from lipid droplets (LDp) and rosette leaves (TOp).

### Arabidopsis caleosins contain putative AIMs and interact with ATG8 proteins

The ATG8 interactors typically possess the AIM/LIR motif with the consensus sequence W/F/Y-X-X-L/I/V (Noda et al., 2010; Liu et al., 2021). As caleosins may potentially mediate LD degradation *via* autophagy, we were interested if they possess putative AIMs. Therefore, by using the iLIR software (Kalvari et al., 2014) we analysed the amino acid sequences of the eight Arabidopsis caleosins and we found that each of them contains at least one putative AIM (Supplemental Table S1, Supplemental Figure S7). The AIM1 (YXXL), localized just before the proline knot, was identified only in CLO1, CLO2, CLO3, and CLO8, whereas the AIM2 (WXXL) was found at the C-termini of all analysed caleosins, except CLO8, which is truncated compared to the others. To verify whether caleosins directly bind to ATG8, we first examined their interaction by using the mating-based split-ubiquitin yeast system. In this approach, we analysed the ATG8b-binding properties of CLO1, CLO2 and CLO3 as well as OLE1, as a positive control. NubWT was used as a positive control to demonstrate that Cub fusion constructs are properly expressed in the yeast cells (Supplementary Figure S8). As shown in Figure 7A and 7B, all tested proteins interacted with ATG8b. A quantitative β-galactosidase assay also revealed that the interaction with ATG8b was comparable for CLO1, CLO3 and OLE1, whereas 2-fold weaker β-galactosidase activity was found for CLO2 (Figure 7C). When ATG8h was used as a prey, detected β-galactosidase activities were lower compared to ATG8b (Figure 7C).

**Figure 7.**
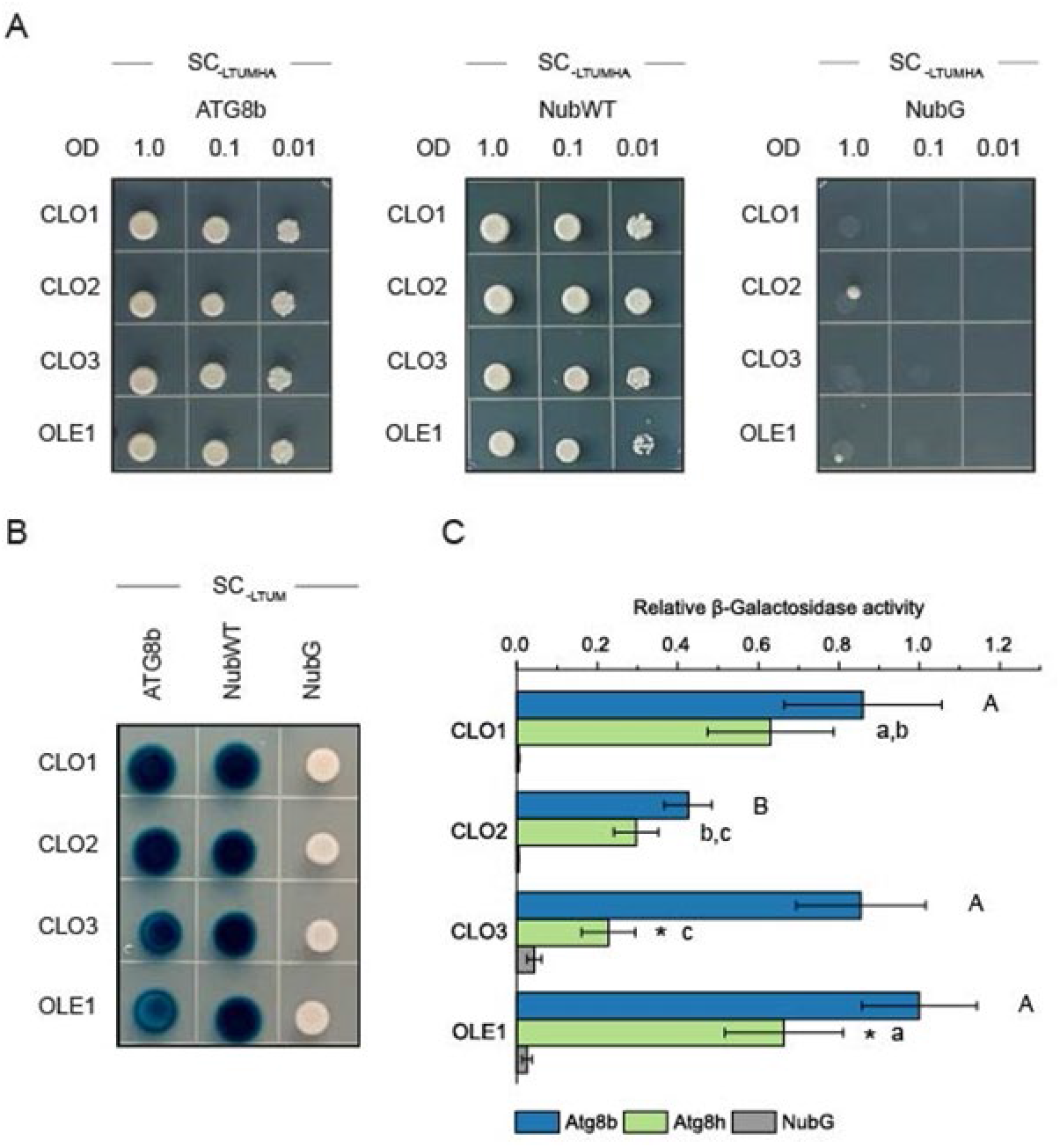
Interaction between caleosins and ATG8 proteins. (A) Yeast mating-based split-ubiquitin assay for interaction of the Cub fusions of CLO1, CLO2, CLO3 or OLE1 (bait) with the NubG fusion of Atg8b (prey). Diploid yeast carrying the given bait and prey constructs were spotted on SC medium without Leu, Trp, Ura, Met, Ade, and His (SC-LTUMAH). Serial dilutions of yeast culture are as indicated. (B) X-Gal overlay assay for detection of β-galactosidase activity for the same bait and prey combinations as in A. Diploid yeast were spotted on SC medium without Leu, Trp, Ura and Met (SC-LTUM). For A and B, NubG and NubI (NubWT) served as a negative and a positive control, respectively. As an additional positive control, interaction between OLE1 and ATG8 proteins was used. (C) Quantitative β-galactosidase activity assay for the Cub fusions of CLO1, CLO2, CLO3 or OLE1 (bait) with the NubG fusions of Atg8b or Atg8h (prey). NubG was used a negative control. Data are means ± SD from 4 (caleosins) or 7-8 (OLE1) independent yeast transformants. The β-galactosidase activity was normalized relative to the activity measured for the interaction between OLE1 and Atg8b. Statistical analysis was performed by one-way ANOVA with Tukey‘s post hoc test and unpaired two-tailed Student’s t test. Different letters indicate significant (Tukey’s test, P < 0.01) differences between baits’ interactions with Atg8b (uppercase) or Atg8h (lowercase). An asterisk denotes significant differences (unpaired two-tailed Student’s t test, P < 0.01) between interaction with Atg8b and interaction with Atg8h for the same bait.

To test, if AIMs are required for interaction of CLO1 with ATG8b, we generated several CLO1 AIM mutants: CLO1-AIM1 (112-117 aa) with deletion of the entire AIM1, CLO1-AIM2 (196-202 aa) with deletion of the entire AIM2, and CLO1-AIM1/AIM2 with deletion of both AIM1 and AIM2. For this step, we have selected the AIM with the highest PSSM score (AIM2, Supplemental Table S1), excluding the ones with a PSSM score below 8 (Supplemental Table S2), except the AIM corresponding to xLIR in CLO3 (AIM1). Additionally, we also generated two CLO1 mutants with deleted N-(1-97 aa, including EF-domain) and C-termini (196-246 aa, including AIM2) (Figure 8A). We found that CLO1-AIM1 and CLO1-AIM1/AIM2 significantly lost their ability to interact with ATG8b as we observed 90% decrease in β-galactosidase activity (Figure 8B). Significant but less prominent decrease in the binding efficiency to ATG8b was also observed for CLO1-AIM2. The proper expression of Cub fusion constructs was assessed by measuring the interaction with NubWT (Figure 8C).

**Figure 8.**
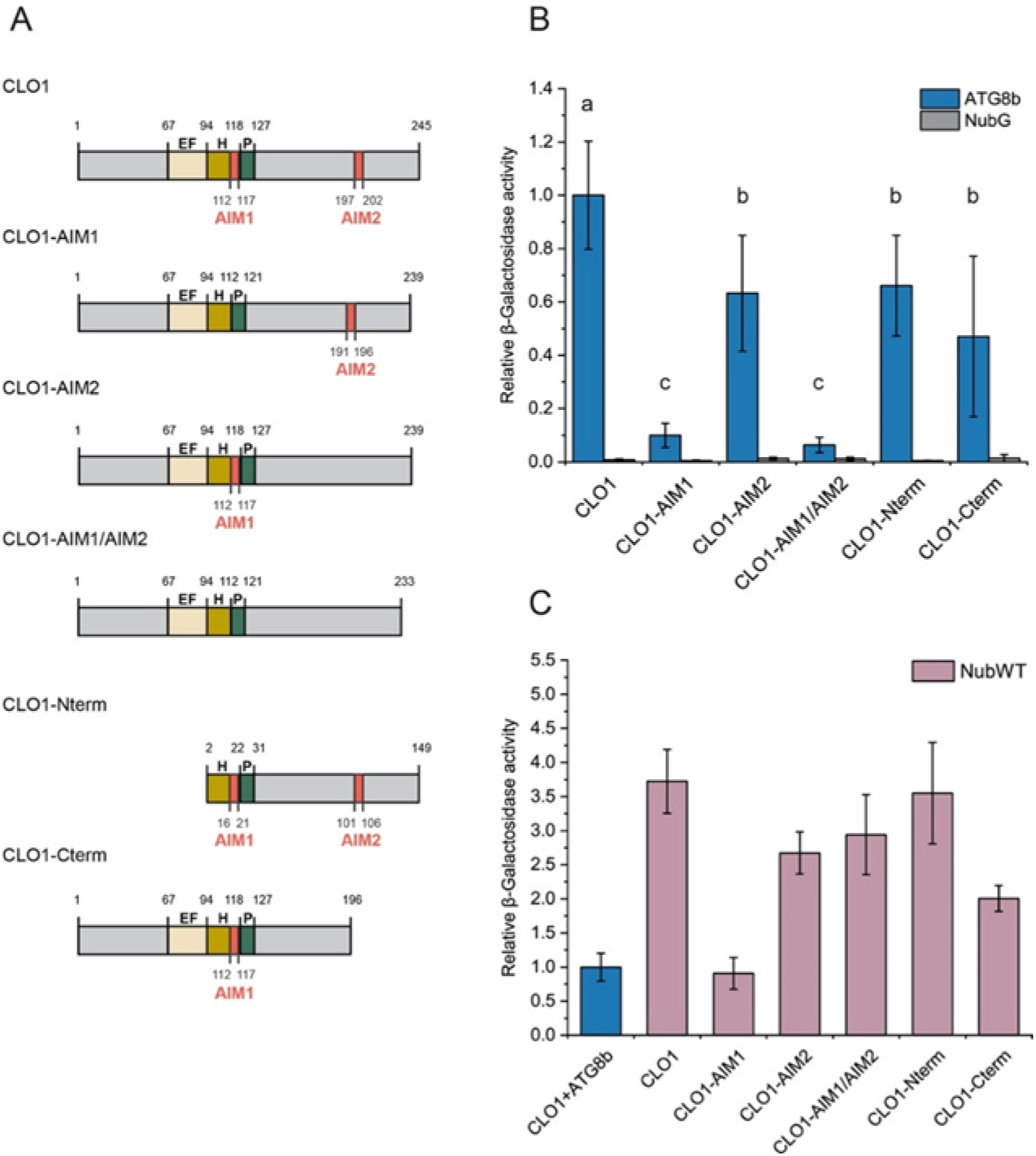
Identification of caleosin motifs mediating interaction with ATG8. (A) Schematic diagram of tested CLO1 variants, lacking AIM1 (112-117 aa; CLO1-AIM1), AIM2 (196-202 aa; CLO1-AIM2), both AIM1 and AIM2 (CLO1-AIM1/AIM2), N-terminus (1-97 aa, including EF-domain; CLO1-Nterm) and C-terminus (196-246 aa, including AIM2; CLO1-Cterm). The EF-hand domain, the central helix and the proline knot are denoted as EF, H and P, respectively. (B) Quantitative β-galactosidase activity assay Cub fusions of CLO1 variants shown in A (bait) with the NubG fusion of ATG8b (prey). NubG was used a negative control. Data are means ± SD from 6-20 (for ATG8b) or 4-6 (for NubG) independent yeast transformants. The β-galactosidase activity was normalized relative to the activity measured for the interaction between CLO1 and ATG8b. Statistical analysis was performed by one-way ANOVA with Tukey‘s post hoc test. Different letters indicate significant (P < 0.01) differences between β-galactosidase activity for the tested variants. (C) Quantitative β-galactosidase activity assay for the Cub fusions of CLO1 variants shown in A with NubWT (positive control). Data are means ± SD from 6 (CLO1) or 4 (CLO1 variants) independent yeast transformants. The β-galactosidase activity was normalized relative to the activity measured for the interaction between CLO1 and ATG8b (blue bar).

As the secondary approach to confirm the binding of CLO1 to ATG8, we performed co-immunoprecipitation assays using the anti-CLO1 antibody. We found that ATG8 co-immunoprecipitated with CLO1 from the LD protein extracts from imbibed seeds of Col-0 but not of *clo1* and *clo1 clo2* mutants. Similar results were also observed during Col-0 seed germination course (Supplementary Figure S9). Together, these data strongly suggest that CLO1 and CLO2 interact with ATG8 proteins and that the putative AIMs identified in CLO1 can be involved in this interaction.

## DISCUSSION

In this study, we have shown that both CLO1 and CLO2 from *A. thaliana* interact with ATG8s, but only the mutation of *CLO1* results in the inhibition of LDs degradation *via* microlipophagy. Consequently, our results are novel with respect to the finding that during seed germination CLO1 may govern the docking and transfer of LDs into the vacuole *via* its interaction with the core autophagosome protein.

### Loss of function of CLO1 affects seed lipid composition and TAGs degradation rates

Disruption of *CLO1* had significant impact on acyl composition of total lipids in *A. thaliana*, resulting in the increase of 18:1 and lower levels of 20:1 in the mature seeds. 20:1 level was also significantly lower in *clo2* seeds. Similar changes in lipid composition in *clo1* and *clo2* were reported recently by Liu et al. (2022) and linked to the function of both caleosins in lipid biosynthesis during *A. thaliana* seed development (Liu et al., 2022). Moreover, *in silico* analysis of Arabidopsis caleosins showed the presence of RY and RAV1 motifs in their promoters which can bind transcription factors regulating oil synthesis during seed development (Shen et al., 2014). Interestingly, suppression of the *OLE1* gene in Arabidopsis had an opposite effect on 18:1 and 20:1 content in mature seeds (Siloto et al., 2006). These findings suggest that core proteins of LDs, besides their structural function, also directly regulate acyl composition of TAGs that accumulate in LDs during the embryo development. Moreover, previous studies on *clo1* mutants demonstrated slower degradation of TAGs in seeds of the *clo1* mutant between 48 h and 60 h of germination, which was identified by elevated levels of 20:1 when compared to wild type seedlings (Poxleitner et al. 2006). We confirmed such a profile of 20:1 in both *clo1* and *clo1 clo2* seeds during later steps of germination (48 h – 96 h). It is thus probable that CLO1 may directly be involved in TAG degradation in germinating seeds and that this pathway functions in addition to the LDs breakdown mediated by SDP1 TAG lipase (Kelly et al., 2011). Notably, despite *sdp1 sdp1L* seeds showed highly reduced TAG breakdown during germination, they were able to germinate, however, with a retarded seedling growth phenotype (Kelly et al., 2011). Therefore, it is possible that when the major lipolysis pathway is inhibited, microlipophagy may be one of the mechanisms facilitating partial mobilisation of storage lipids and in consequence seed germination, albeit at limited rates.

### No trafficking of LDs to the vacuoles is observed in *clo1* during seed germination

Our observations of Col-0 germinating seeds overexpressing the tonoplast marker protein and labelled for LDs showed that individual LDs found in vacuoles often associate with the tonoplast invaginations. We found also that during germination of both Col-0 and *clo2* seeds, LDs appeared in the central vacuoles together with the cytoplasmic material, which is characteristic for microlipophagy. Consequently, the later steps of seed germination were accompanied by the presence of LDs undergoing degradation inside of the vacuole in these plants. In contrast, in *clo1* and *clo1 clo2* mutants, no LDs were found in the vacuoles. Instead, they accumulated at the peripheral parts, of the cells forming aggregates around the vacuoles. Similar observations have been reported previously for *clo1* plants by Poxleitner et al. (2006). Such distribution of LDs was particularly visible during germination of *clo1* and *clo1 clo2* seeds under continuous dark conditions. These results indicate that loss of function of CLO1, but not of CLO2, inhibits the transfer of LDs to the vacuoles and in consequence their degradation *via* microlipophagy pathway. To date, most of reports have described the role of microlipophagy in LDs degradation only in non-seed tissues (for review see: Zienkiewicz and Zienkiewicz (2020); Xu and Fan (2022)). For example, Fan et al. (2019) showed that LDs found inside the vacuole or within invaginations of the tonoplast in dark-treated *A. thaliana* rosette leaves undergo degradation *via* microlipophagy-resembling process. LDs breakdown by microlipophagy has been also recently demonstrated in developing and germinating pollen of angiosperms (Zhao et al., 2020; Akita et al., 2021). Overall, our findings show that in germinating seeds a certain pool of LDs is transported to the vacuoles by tonoplast invaginations and for their degradation and we suggest that microlipophagy is rather universal process occurring in different plant tissues.

### ATG8 co-localize with LDs

In this study, the autophagic marker, protein ATG8b, was found to co-localize with LDs during the course of seed germination. Nevertheless, this co-localization was restricted to a limited pool of LDs and intensified under continuous dark conditions. After determining localization of LDs inside of the vacuoles by electron microscopy, we did not however observe an association of macroautophagic membrane structures around the LDs. Similar pattern of co-localization was also reported for DsRed-ATG8e-labelled puncta and LDs in Arabidopsis leaves under dark-induced starvation by Fan et al. (2019). Moreover, in the same study no macroautophagic membrane structures were found around LDs localized inside of vacuoles and therefore suggested that in Arabidopsis leaves degradation of LDs occurs *via* microlipophagy pathway. Interestingly, other study on Arabidopsis leaves showed that GFP-ATG8-labelled structures localized at photodamaged chloroplasts but they never completely surrounded the entire chloroplast (Nakamura et al., 2018). Based on their results, the authors proposed that ATG8-associated structures are essential for triggering the chlorophagy and are also directly involved in the tonoplast-mediated sequestering of chloroplasts (Nakamura et al., 2018). We propose that similar ATG8-depending mechanisms govern the LDs trafficking between the cytoplasm and the vacuoles prior to their degradation during seed germination.

Our immunoblotting studies showed that during seed germination ATG8-PE conjugates are abundantly present in the protein fraction isolated from LDs, indicating that ATG8-PE is indeed associated with LDs. Accumulation of the lipidated ATG8 form has also been reported for Arabidopsis leaves under darkness-induced starvation (Fan et al., 2019). In the same study it was also found that degradation of LDs *via* microlipophagy-resembling process relies on the core ATG proteins, including ATG2 and ATG5 (Fan et al., 2019). Interestingly, the latter protein is crucial for lipidation of ATG8 (Martens and Fracchiolla, 2020) as well as for maintaining TAG degradation, as shown in *atg5-1* and *atg5-3* mutans of Arabidopsis (Avin-Wittenberg et al., 2015).

### ATG8 interacts with structural proteins of LDs

To investigate which of the LD structural proteins could directly interact with ATG8s, we searched for AIMs in *A. thaliana* caleosins and OLE1. The latter protein was previously shown to interact with ATG8 by a LIR/AIM docking site described by Marshall et al. (2019). Interestingly, as AIMs with position-specific scoring matrix (PSSM) higher than 13 are considered to be reliable predictions (Kalvari et al., 2014; Klionsky et al., 2016), two WxxL motifs with a very low PSSM value identified in the OLE1 sequence by the iLIR tool are most likely not functional AIMs. However, it should be noted that numerous ATG8-interacting plant proteins that contain AIMs with a PSSM score below 13 have also been identified (Liu et al., 2021). Furthermore, similar to Marshall et al. (2019), we confirmed binding of OLE1 to ATG8b and ATG8h proteins in our split-ubiquitin two-hybrid assays. These data suggest that interaction between OLE1 and ATG8 may occurs *via* AIM yet to be determined using other algorithms.

Interestingly, we found putative AIMs (AIM1/AIM2) with relatively high PSSM scores (between 7 and 17) in all analysed caleosins. We also confirmed that CLO1, CLO2 and CLO3 interact with ATG8 proteins. The stronger interaction between CLO1 and ATG8b/ATG8h compared to CLO2 may explain the loss of microlipophagy phenotype that was observed only in *clo1* mutants. Notably, the interaction between ATG8b and CLO3 suggests that this leaf-specific caleosin may be important for regulating microlipophagy in Arabidopsis leaves.

To gain more insights into the role of AIMs in the binding of caleosin to ATG8, we generated several CLO1 mutants with deletions in both AIM1 and AIM2 motifs. While the deletion of the C-terminal AIM2 caused a slight decrease in the binding of CLO1 to ATG8b, the absence of the transmembrane-localized AIM1 almost completely abolished the interaction between the two proteins, suggesting that AIM1 plays a prominent role in binding of CLO1 to ATG8b. To date, several functional AIMs in transmembrane regions have been reported in diverse plant proteins (Stephani and Dagdas, 2020; Zhang et al., 2020). Similarly to AIMs in maize reticulon proteins Rtn1/2 (Zhang et al., 2020), AIM1 in caleosins seems to be located close to the cytoplasmic face of the LDs lipid monolayer and therefore can be partly accessible to ATG8 proteins, even when caleosins are still embedded in the LDs membrane. Impaired binding to ATG8 observed for N- and C-terminal truncated mutants of CLO1, despite of the presence of the full length AIM1, might be the result of structural changes of the protein affecting CLO1 targeting and its orientation at the LDs membrane. Indeed, N-terminal truncated mutants of Arabidopsis CLO1 (up to 95 amino acids) (Purkrtová et al., 2015) as well as sesame seed caleosin, structurally similar to *At*CLO1, which lacks the helical subdomain (residues 100-115) (Chen and Tzen, 2001) have been shown to possess impaired targeting to LDs. Alternatively, the interaction of caleosins with ATG8 can also be mediated by other motifs, such as the ubiquitin-interacting motif (UIM) (Marshall et al., 2019) or *via* not yet discovered mechanisms. Noticeably, all deletion mutants of CLO1 tested in our study interacted with NubWT, indicating that they were properly synthesized and stable in yeast cells.

In summary, our work demonstrates that CLO1 functions as a mediator in trafficking of LDs to the vacuole for their degradation. In addition, we provide evidence that ATG8 localizes to LDs *via* the interaction with CLO1. As illustrated in Figure 9, we propose that the docking of ATG8 to CLO1 may promote structural interaction and fusion of LDs with the tonoplast, followed by the transfer of LDs into the vacuole, where the degradation process takes place (Figure 9A). Alternatively, interaction between ATG8 and CLO1 may trigger degradation of LDs structural proteins enabling the fusion between the membrane of LDs and the tonoplast (Figure 9B). Our data also revealed that the interaction of OLE1 with ATG8 is not sufficient to trigger the trafficking of LDs into the vacuoles in Arabidopsis seeds. Although further studies will be required to determine the molecular mechanisms behind the interaction of LDs with the vacuoles in plant cells, the crucial role of a specific caleosin in this process is now firmly established in germinating seeds. Consequently, our findings open new avenues of research exploring integrated aspects of lipid mobilization in diverse plant tissues.

**Figure 9.**
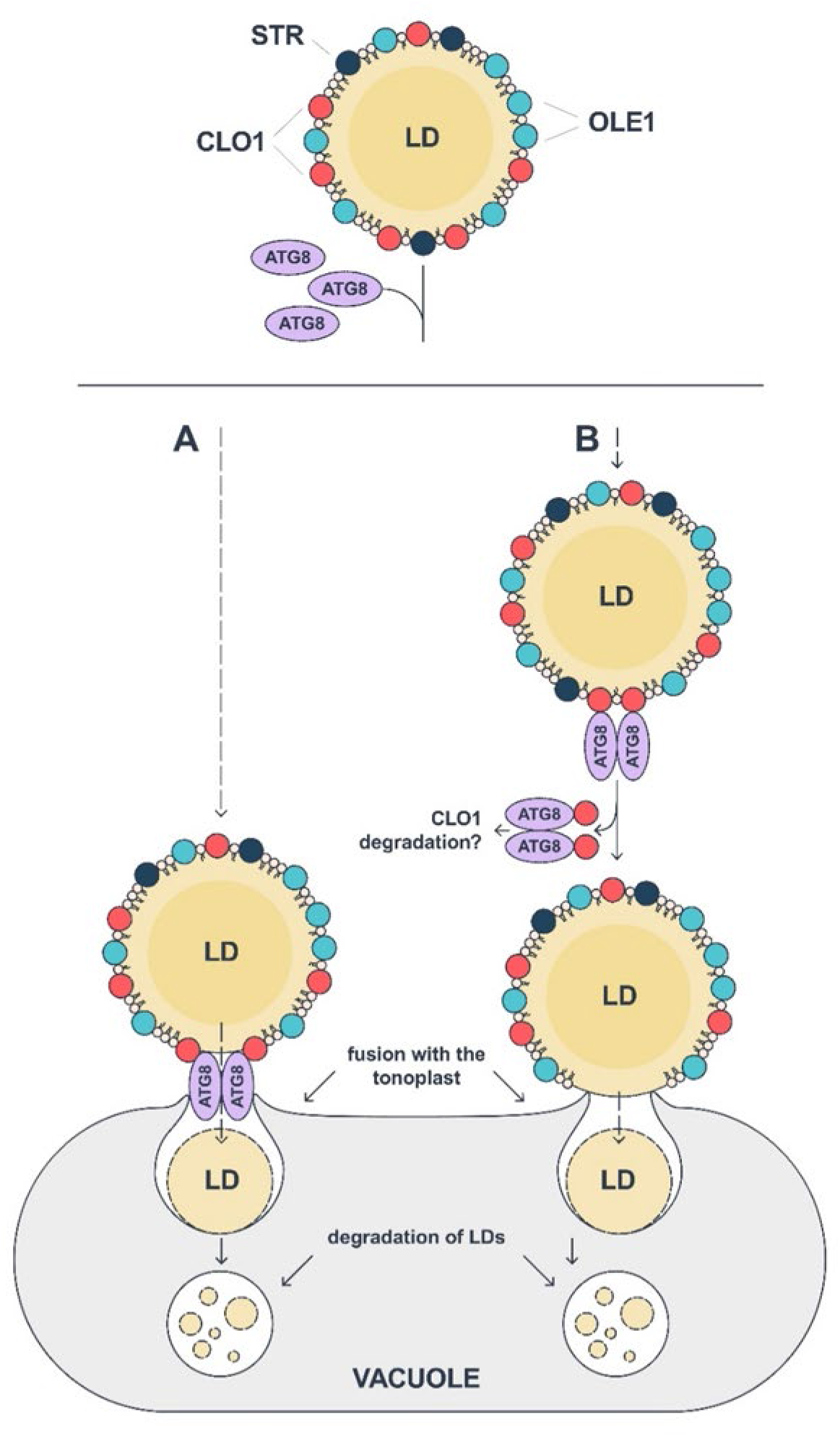
Model of CLO1-mediated microlipophagy in Arabidopsis seeds. Interaction of ATG8 with CLO1 triggers the fusion of LDs with the tonoplast *via* two possible scenarios: (A) the complex of CLO1 and ATG8 directly mediates the fusion of the LD membrane with the tonoplast, and (B) following its binding to CLO1, ATG8 promotes degradation of CLO1 resulting in direct interaction and fusion between LDs and vacuole membranes. Once transported to the vacuole, LDs undergo a gradual degradation during seed germination. CLO1, caleosin 1, OLE1, oleosin 1; LD, lipid droplet, STR; steroleosin.

## MATERIALS AND METHODS

### Plant materials and growth conditions

The seeds of *Arabidopsis thaliana* T-DNA insertion lines for CLO1 (*Atclo1-1*, Poxleitner et al., 2006) and CLO2 (SALK_046559) in the Columbia 0 (Col-0) accession were obtained from the Nottingham Arabidopsis Stock Centre (NASC). The Col-0 ecotype was used as a wild type control. The *clo1* mutant was previously described by Poxleitner et al. (2006). Homozygous lines were identified by PCR using the left border primer of the T-DNA insertion and the allele-specific primers listed in Supplemental Table S3. *A. thaliana* lines containing indicator for the tonoplast - γ-TIP, generated with cyan fluorescent protein (vac:CFP), were obtained from NASC under NASC stock number: N16256. For soil grown plants, seeds were sown on soil, cold-stratified for 48 h and transferred to a growth chamber (22°C, 16 h light/8 h dark, 120– 150 µmol m^-2^ s^-1^). For plate grown plants, seeds were surface-sterilized with 30% (v/v) bleach solution, washed four times with distilled water and placed on agar-solidified half strength Murashige and Skoog (MS) medium. After stratification at 4°C in the dark for 2 days, the plates were incubated at 22°C under a 16 h light/8 h dark photoperiod, continuous dark (24 h of dark) or continuous light (24 h of light).

### Seed phenotypic analysis

Mature seeds were collected and then photographed using a Olympus SZX12 stereomicroscope with R6 Retiga digital camera (Qimaging) Length and width measurements were performed using ImageJ software (Schneider et al., 2012).

### Total RNA extraction and cDNA synthesis

Total RNA was extracted from 100 mg of *A. thaliana* siliques, germinating seeds or rosette leaves using Spectrum™ Plant Total RNA Kit (Merck KgaA, Darmstadt, Germany) according to the manufacturer’s instructions. Total RNA (1 µg) was used for cDNA synthesis using RevertAid H Minus First Strand cDNA Synthesis Kit and Oligo(dT)18 primer (Thermo Fisher Scientific, Bremen, Germany).

### Construction of plasmids and Arabidopsis transformation

The coding sequence for *CLO1* and *ATG8b* were amplified from cDNA using Phusion High-Fidelity DNA polymerase (Thermo Fischer Scientific) and attB primers listed in Supplemental Table S4. The amplified sequences were inserted into the pDONR207 vector and then transferred into the pEarleyGate 104 (YFP at the N-terminus) vector using Gateway technology (Earley et al., 2006). The resulting plasmids were introduced to *Agrobacterium tumefaciens* strain GV3101 and used for transformation of *A. thaliana via* floral dipping (Clough and Bent, 1998). T1 transgenic lines were selected by spraying 7-day old seedlings with 0.05% Basta solution and surviving plants were subjected to Western blot analysis for the detection of EYFP (see below). T3 transgenic lines were used for microscopic analyses (see below).

### Total protein isolation

Plant material was homogenized with a TissueLyser II (Qiagen) and protein extraction buffer (100 mM Tris pH 8.0, 2 mM phenylmethylsulfonyl fluoride (PMSF), 2% (v/v) β-mercaptoethanol, 4% (w/v) SDS), was added (0.5 ml) to the samples. The samples were then heated for 3 min at 80°C and centrifuged at 13,000*g* for 5 min. The supernatant was transferred to a new tube and used for SDS-PAGE.

### Isolation of LDs and LD-associated protein

LDs were isolated as described in Rudolph et al. (2011) with modifications. Briefly, mature and/or germinating seeds (24 h and 48 h) of *A. thaliana* (Col-0, *clo1*, *clo2*, *clo1clo2*) were homogenized at 4°C in grinding buffer (15% (w/v) sucrose, 150 mM TRIS-HCl, pH 7.5, 10 mM KCl, 1.5 mM EDTA, 0.1 mM MgCl2, and 5 mM dithiothreitol). The homogenates were centrifuged at 100,000 *g* for 60 min at 4°C. The floating fat pad was collected and resuspended in cold flotation buffer (10% (w/v) sucrose, 50 mM TRIS-HCl buffer, pH 7.5, 10 mM KCl, 1.5 mM EDTA, and 0.1 mM MgCl2). After centrifugation at 100,000*g* for 60 min at 4°C, the resulting fat pad was collected and homogenized in 0.2 M borate buffer, pH 8.25. Proteins were precipitated at -20°C, overnight using acetone/ethanol (4:1, v/v), followed by washing with 80% ethanol.

### SDS-PAGE and Immunoblotting

Protein samples (25 µg) were mixed with 2 x Laemmli sample buffer and boiled for 5 min. Then, proteins were subjected to 12% SDS-PAGE and transferred onto nitrocellulose membrane. Membranes were incubated for 1 h in blocking solution containing 1% (w/v) non-fat dry milk in TRIS-buffered saline plus Tween 20 (TBST, 20 mM Tris-HCl pH 7.5, 150 mM NaCl and 0.05% (v/v) Tween 20, pH 7.5). Membranes were subjected to immunoblot analysis using primary antibodies against CLO1 [diluted 1:1000, (Næsted et al., 2000)], OLE1 [diluted 1:1000, (Rudolph et al., 2011)], ATG8 (diluted 1:500, catalog no. AS 142769, Agrisera) or GFP (diluted 1:1000, catalog no. 11814460001, Roche). Alkaline phosphatase conjugated anti-rabbit, anti-mouse or anti-chicken IgG (diluted 1:30 000, Merck), respectively, served as a secondary antibody. Proteins were visualized on nitrocellulose membrane as a purple-brown bands in the presence of nitrotetrazolium blue chloride (NBT) and 5-bromo-4-chloro-3’-indolyphosphate p-toluidine (BCIP).

### Co-IP Assay

LD-associated proteins were co-immunoprecipitated using the Dynabeads™ Co-immunoprecipitation Kit (Thermo Fisher Scientific) and an anti-CLO1 antibody according to the manufacturer’s protocol for Western Blot. The obtained Dynabeads Co-IP complexes were processed according to the manufacturer’s protocol. Proteins bound to Dynabeads were eluted with 40 µl of elution buffer and analysed by SDS-PAGE. For immunoblotting, the membrane was cut between the 25 kDa and the 15 kDa marker bands and the upper part of the membrane was probed with the anti-CLO1 antibody while the lower part was probed with the anti-ATG8 antibody as described above.

### Yeast split-ubiquitin assay

The direct protein-protein interactions were analysed using the mating-based split-ubiquitin system (mbSUS) (Obrdlik et al., 2004; Grefen et al., 2007). The coding regions of the selected caleosin and oleosin genes were amplified using Phusion High-Fidelity DNA polymerase (Thermo Fischer Scientific) and primers listed in Supplemental Table S5. Site-directed mutagenesis of ATG8-Interacting Motifs (AIM) was performed using Q5 site-directed mutagenesis kit (New England BioLabs, Frankfurt am Main, Germany) using primers listed in Supplemental Table S5 according to the manufacturer’s protocol. AIMs were identified in caleosin sequences and OLE1 using iLIR tool (http://repeat.biol.ucy.ac.cy/iLIR/, Kalvari et al. (2014)). PCR products were inserted into pMetYCgate or pNXgate32-3HA vectors by *in vivo* recombination in THY.AP4 and THY.AP5 yeast strains, respectively, as described in Grefen et al. (2007). Transformants were selected on synthetic complete (SC) media lacking leucine (THY.AP4) or tryptophan (THY.AP5). The bait or prey vectors were isolated from several yeast clone cultures, transformed into *Escherichia coli* for re-isolation and sequencing. After sequence confirmation, the bait or prey vectors were again transformed into THY.AP4 and THY.AP5 strains, respectively. The obtained colonies were subsequently used for mating in appropriate combinations. Diploids were selected by replica plating on SC medium lacking leucine, tryptophan, uracil and methionine. In order to detect the protein-protein interactions, diploid colonies were grown on liquid SC medium lacking leucine, tryptophan, uracil and methionine overnight at 28° C. To confirm the interactions, yeast cells (100 μl) were harvested and diluted in sterile water to an OD600 of 1.0, 0.1 and 0.01 and 4 μl of each dilution was spotted onto a selective medium and grown for 1 to 3 days at 28°C. The remaining yeast cells were used for a total protein extraction. Total yeast protein extraction was performed according to the protocol described by Horaruang and Zhang (2017). Immunoblot analysis with anti-HA antibody was performed as described above. The selected interactions were verified by quantitative β-galactosidase assays using Beta-Glo Assay System (Promega, Madison, WI, USA). Briefly, 10 μl of the overnight culture diluted to an OD600 of 0.02 was mixed with 10 μl of Beta-Glo® Reagent and the assay was performed according to the manufacturer’s protocols. X-Gal overlay assay was used as an additional method to confirm protein-protein interactions and was carried out as described in Grefen et al. (2007). The soluble wild type NubI (the pNubWT-Xgate vector) and NubG (the empty pNXgate32-3HA vector) were used as a positive and a negative control, respectively. Additionally, pMetYCgate carrying *AtOLE1* was used as a positive control (Marshall et al., 2019).

### Fatty acid analysis

Lipids were extracted from mature seeds and seedlings germinated on MS medium as described by Marmon et al. (2017). Briefly, 1 ml of methanolic solution containing 2.75% (v/v) H2SO4 (95%–97%), 2% (v/v) dimethoxypropan and 2% (v/v) toluene was added to the plant material. Samples were shaken at 80°C for 45 min and washed with 1 ml *n*-hexane and 1.2 ml of saturated NaCl solution. After centrifugation at 1000 g for 10 min, the upper phase was collected and the samples were dried under nitrogen streaming. Fatty acid methyl esters (FAMEs) analysis was performed according to Marmon et al. (2017). For quantification, an internal standard of tripentadecanoate (tri-15:0) was added to each sample before the methylation.

### Confocal laser scanning microscopy (CLSM)

Lipid droplets were observed using BODIPY™ 505/515. Plant material was incubated with 1 mM BODIPY™ 505/515 or for 30 min and washed with phosphate-buffered saline (PBS, pH 7.2). Samples were observed under Leica TCS SP5 or Olympus F3000 confocal microscope with excitation at 505 nm and emission at 515 nm for BODIPY. EYFP fusion proteins were observed with excitation at 513 nm and emission at 527 nm. CFP fusion protein was observed with excitation at 405 nm and emission at 485 nm. Chlorophyll autofluorescence was captured using excitation at 633 and emission at 700 nm. All images were processed using ImageJ software (Schneider et al., 2012).

### Transmission Electron Microscopy (TEM)

Cotyledons were collected after 48 h or 72 h of seed *in vitro* germination. Sample preparation was carried out as described in Zienkiewicz et al. (2020). Briefly, the samples were fixed in 2.5% (v/v) glutaraldehyde in 0.1 M cacodylate buffer (pH 7.2) overnight at 4°C. Next, the samples were postfixed in OsO4 at room temperature for 1 h, dehydrated in ethanol series and embedded in Araldite-Epon resin (Araldite 502/Embed 812 Kit; Electron Microscopy Sciences, Science Services GmbH, München, Germany). Ultrathin sections were observed with a JEOL 1011 transmission electron microscope (Japan Electron Optics Laboratories Germany GmbH, Freising, Germany) at 80 kV. Images were processed using ImageJ software (Schneider et al., 2012).

### Sequence analysis

The protein sequences of *A. thaliana* caleosins with the following accession numbers: CLO1(O81270); CLO2 (Q9FLN9); CLO3 (O22788); CLO4 (Q9CAB7); CLO5 (B3H7A9); CLO6 (Q9CAB8); CLO7 (F4I4P8); CLO8 (A0A178V895) were retrieved from UniProt database (www.uniprot.org) and aligned by ClustalW using MEGA-X software (Kumar et al., 2018). In this paper, we followed the nomenclature of *A. thaliana* caleosins proposed by Shen et al. 2014 (Supplemental Table S6).

### Statistical analyses

Statistical analyses were performed using R 4.2 software and Statistica 13.1 software (StatSoft, Poland). Statistically significant differences were determined by one-way analysis of variance (ANOVA), followed by post-hoc Tukey’s test, or by a two-tailed Student’s t test (Supplemental Data Set 2).

## AUTHOR CONTRIBUTIONS

AZ designed the study; MM, AZ, KZ, MR, KEB performed the experiments; IF provided materials, analysed data and advised on experimental design; MM and AZ collected, analysed or interpreted the data; MM and AZ wrote the manuscript with contributions from all co-authors.

## ACKNOWLEDGEMENTS

We thank Dr. Corinna Thurow (Department of Plant Molecular Biology and Physiology, University of Göttingen, Göttingen, Germany) for her support with the β-galactosidase assays, Prof. Volker Lipka and Dr. Elena Petutschnig from the Central Microscopy Facility of the Faculty of Biology & Psychology (University of Göttingen, Göttingen, Germany) for the possibility to use Leica TCS SP5 CLSM microscope. We are grateful to Prof. Anthony H. C. Huang for providing us with the anti-oleosin antibody and to Prof. Marianne Poxleitner for providing us with the anti-caleosin antibody. We acknowledge Małgorzata Miklaszewska (https://www.linkedin.com/in/miklassic/) for preparing Figure 9.

## TABLES

**Table S1.**
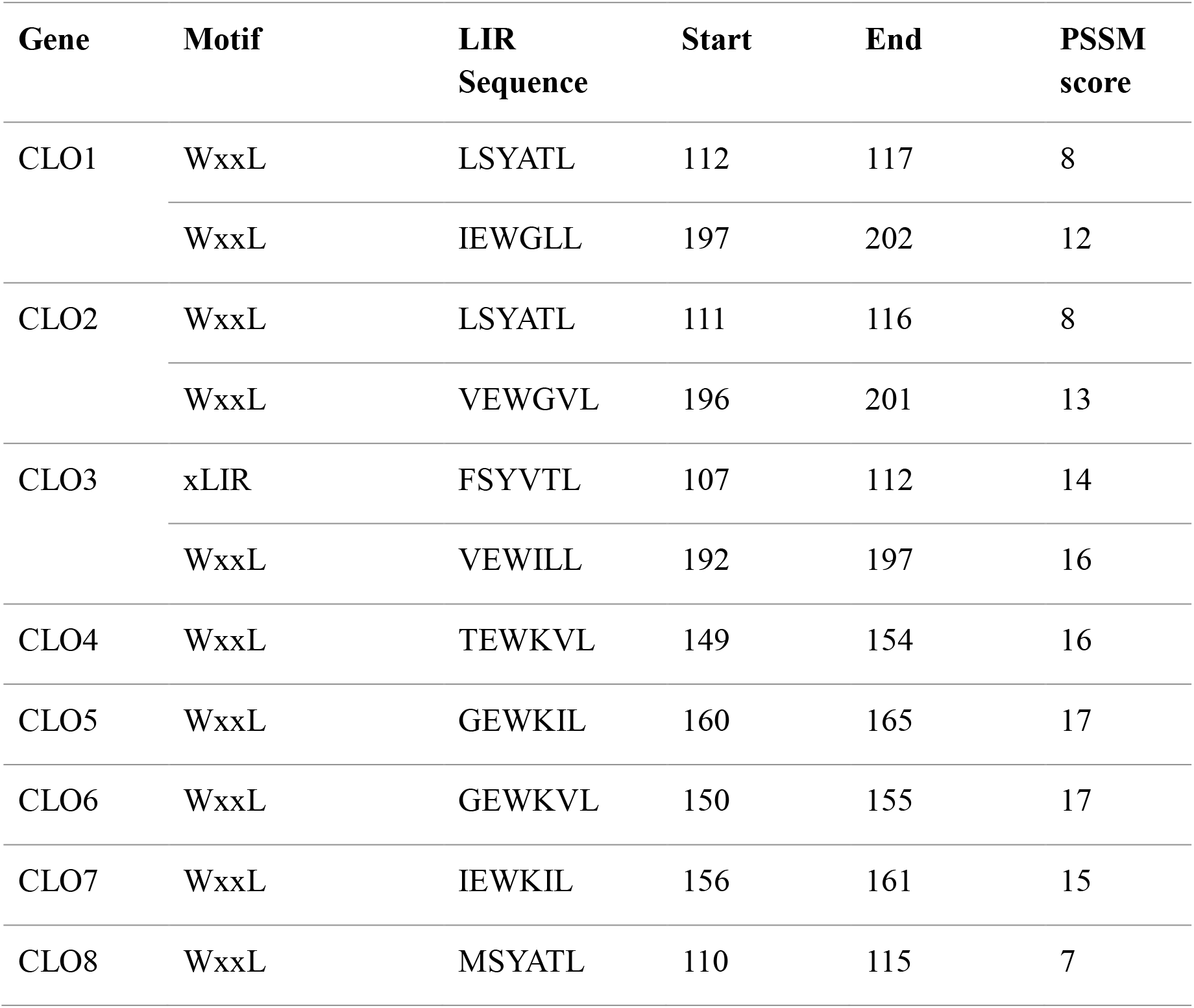
Putative ATG8-Interacting Motifs (AIMs) in caleosins from A. thaliana. Summary of AIMs with a PSSM value above 13 (or corresponding AIMs with a lower PSSM score in other caleosin sequences) identified using the iLIR tool (http://repeat.biol.ucy.ac.cy/iLIR/, Kalvari et al. (2014)); PSSM: position-specific scoring matrix; WxxL: canonical LIR motif; xLIR: extended LIR-motif.

**Table S2.** Putative ATG8-Interacting Motifs (AIMs) with a low PSSM score in CLO1 and OLE1. Summary of AIMs with a PSSM value below 13 identified using the iLIR tool (http://repeat.biol.ucy.ac.cy/iLIR/, Kalvari et al. (2014)); PSSM: position-specific scoring matrix; WxxL: canonical LIR motif.

| **Gene** | **Motif** | **LIR Sequence** | **Start** | **End** | **PSSM score** |
| --- | --- | --- | --- | --- | --- |
| CLO1 | WxxL | APYAPV | 17 | 22 | 4 |
|  | WxxL | VSFFDI | 71 | 76 | 5 |
|  | WxxL | ETYSGL | 86 | 91 | 8 |
|  | WxxL | LGFNII | 94 | 99 | 7 |
|  | WxxL | SPFFPI | 123 | 128 | 5 |
|  | WxxL | GRFMPV | 149 | 154 | 4 |
|  | WxxL | DIFGWI | 188 | 193 | 2 |
|  | WxxL | KIYAGI | 232 | 237 | 2 |
| OLE1 | WxxL | VIFSPI | 79 | 84 | 2 |
|  | WxxL | TVFSWI | 111 | 116 | 0 |

**Table S3.**
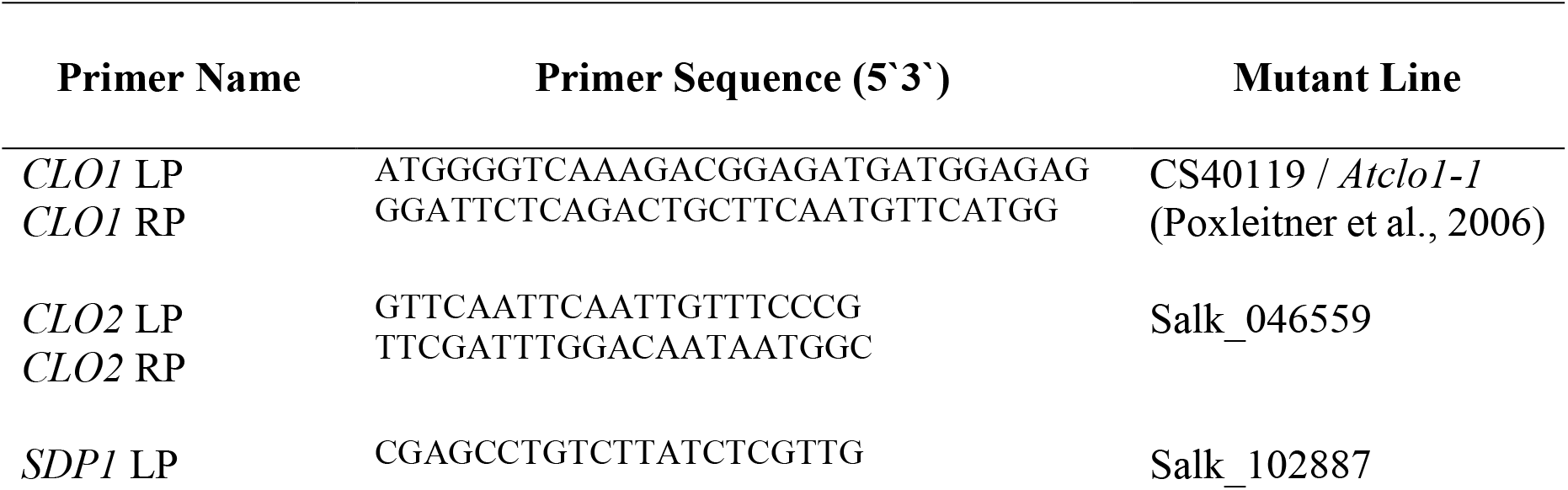
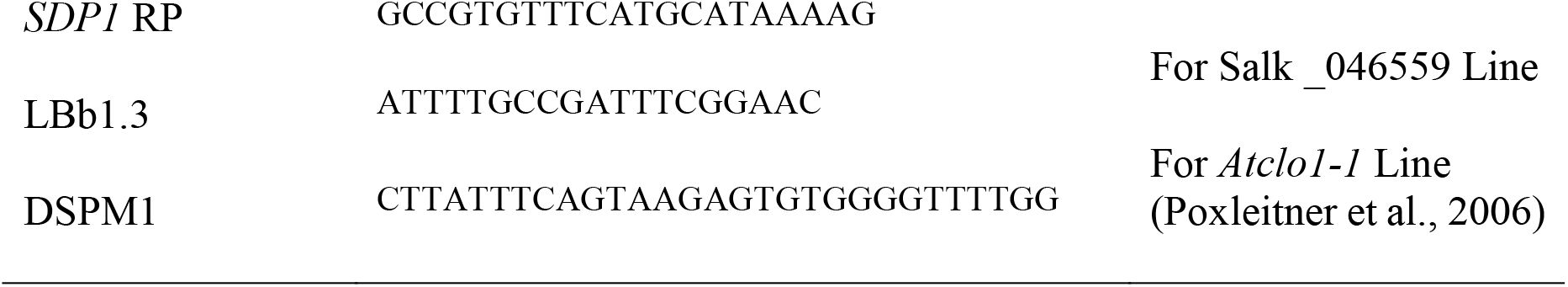
Gene specific primers (LP – left primer, RP-right primer) and T-DNA left border primers (LBb1.3 and DSPM1) used for genotyping of plants.

**Table S4.**
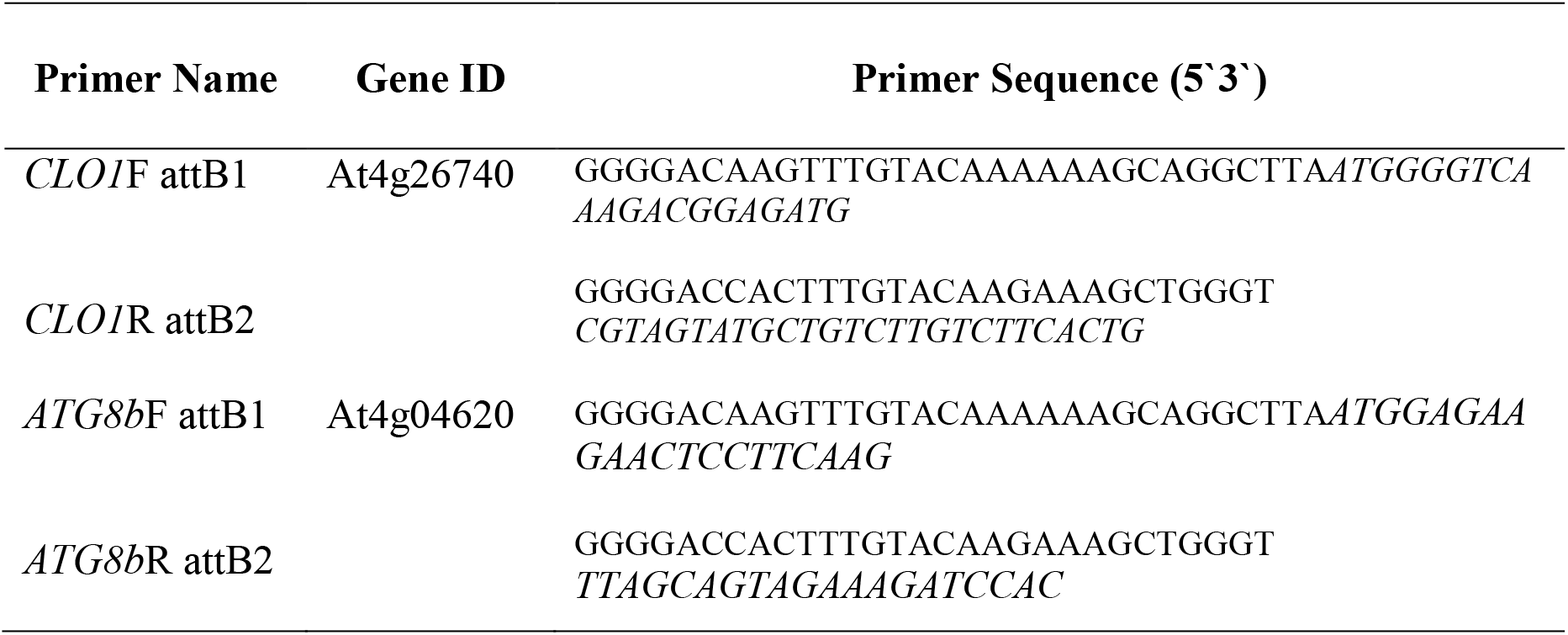
Gene specific primers (F – forward primer, R-reverse primer) used for construction of 35S::EYFP plasmid.

**Table S5.**
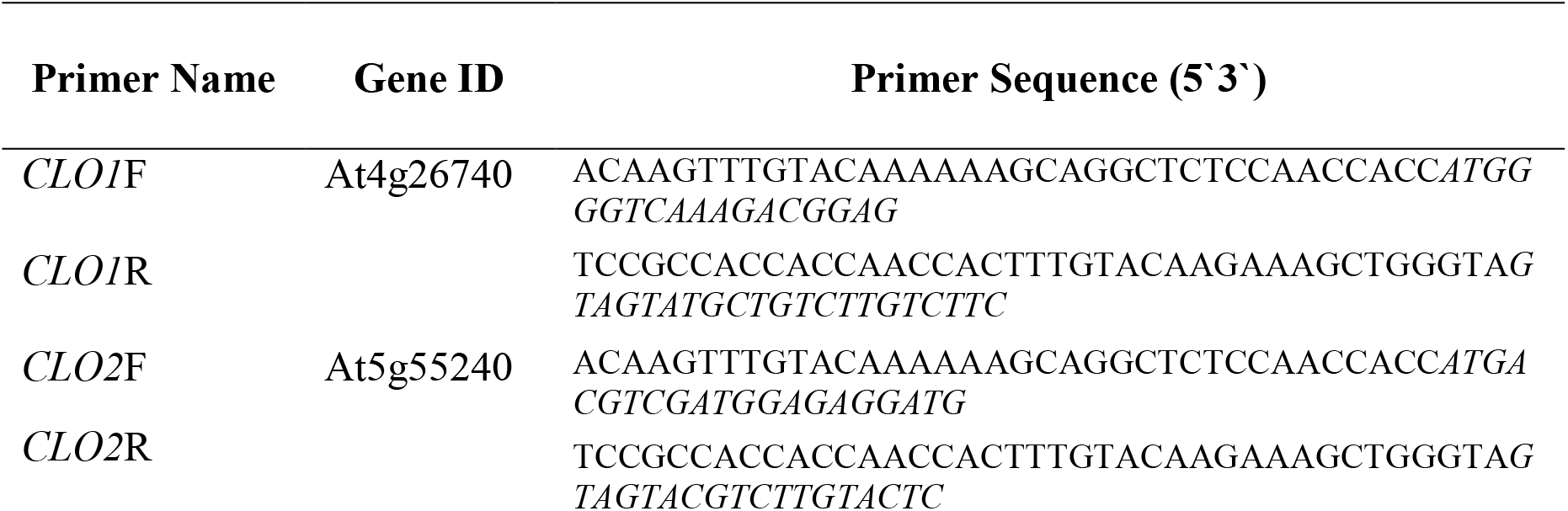
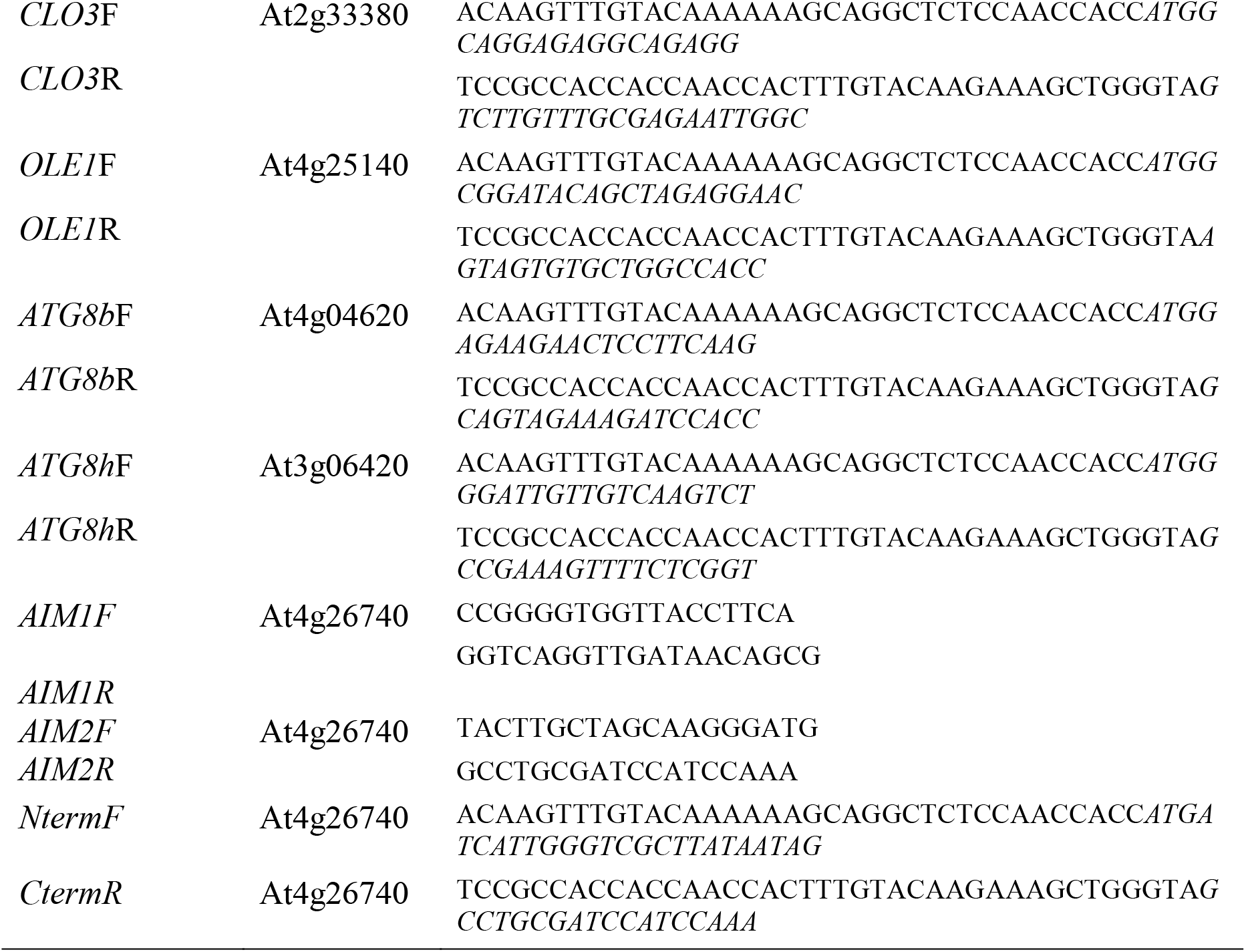
Gene specific primers (F – forward primer, R-reverse primer) used for construction of the yeast plasmids.

**Table S6.**
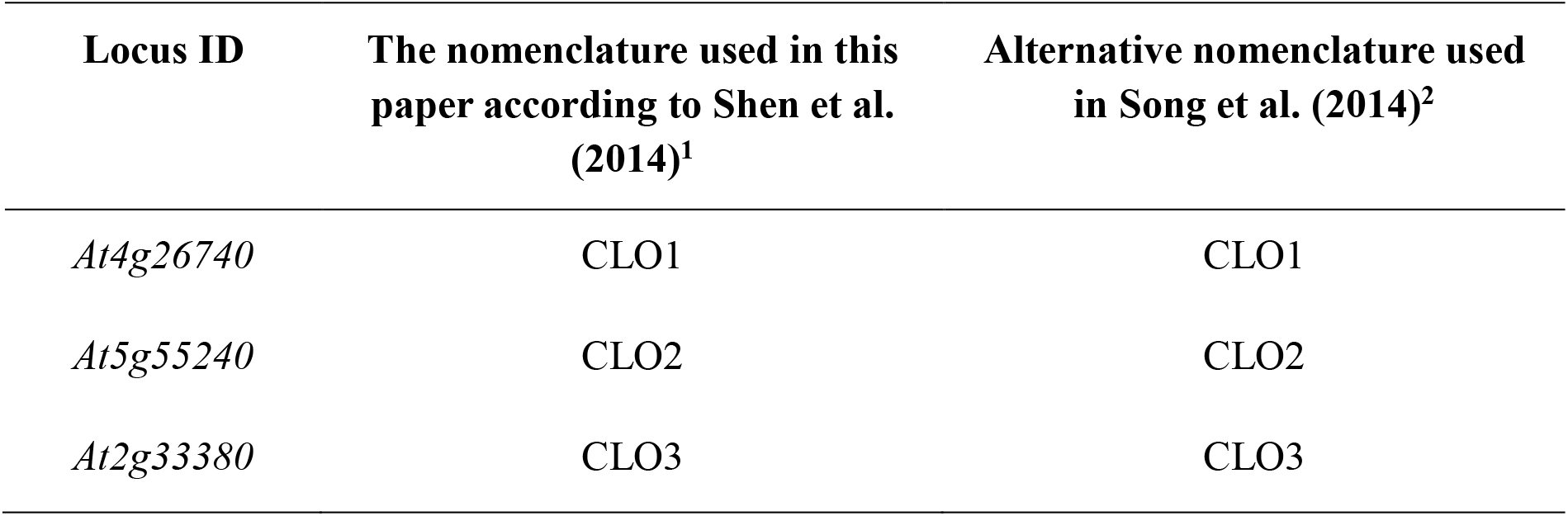
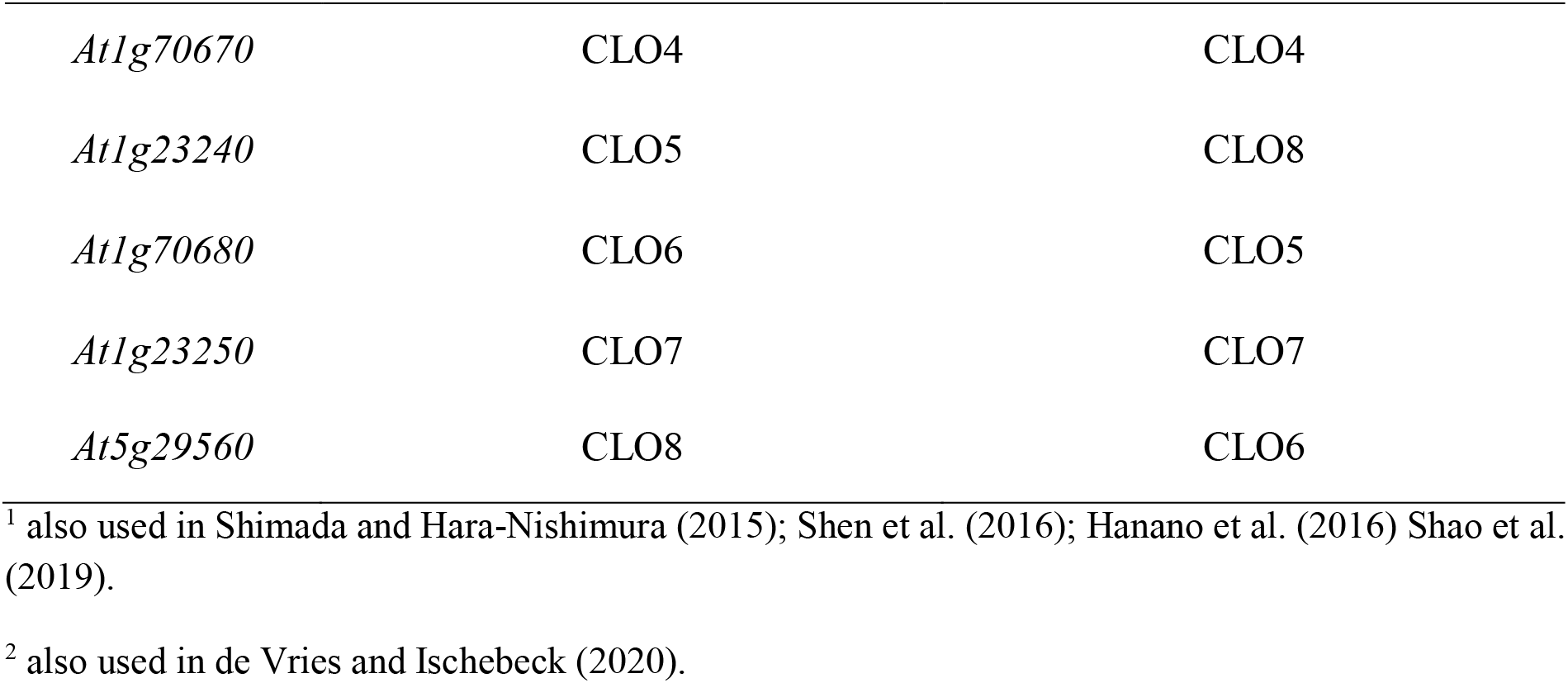
The nomenclature of caleosins from Arabidopsis thaliana used in the literature.

## SUPPLEMENTAL FIGURES

**Supplemental Figure S1.**
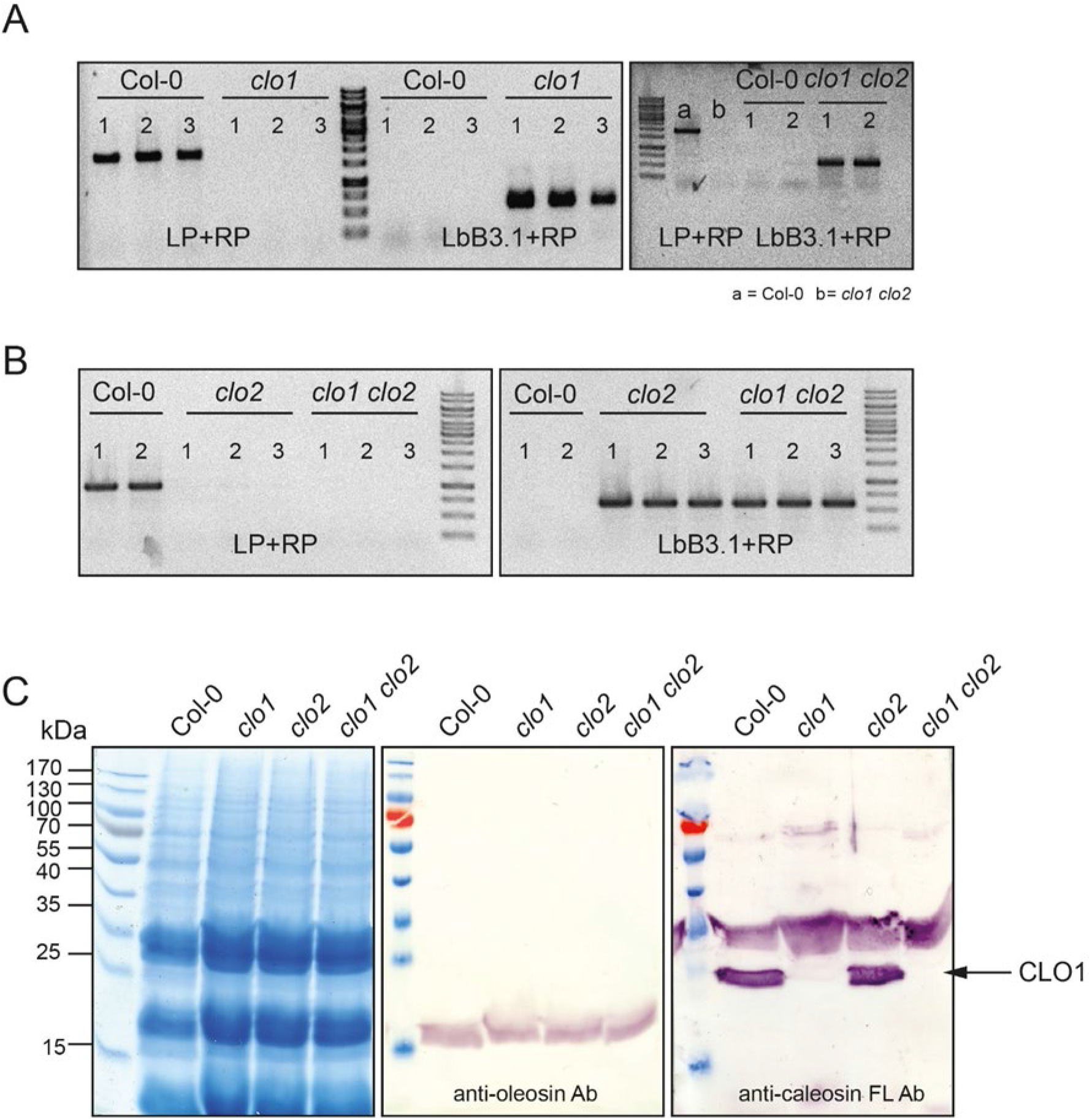
Characterization of T-DNA insertion *clo1* and *clo2* mutant lines of Arabidopsis. (A) Genotyping of *clo1* and *clo1 clo2* mutants by RT-PCR using left border primer of the T-DNA insertion (LbB3.1) and *CLO1* allele-specific primers (LP and RP). (B) Genotyping of *clo2* and *clo1 clo2* mutants by RT-PCR using left border primer of the T-DNA insertion (LbB3.1) and *CLO2* allele-specific primers (LP and RP). (C) Detection of oleosin and caleosin 1 in the total protein extract isolated from Col-0, *clo1*, *clo2* and *clo1 clo2* mature seeds.

**Supplemental Figure S2.**
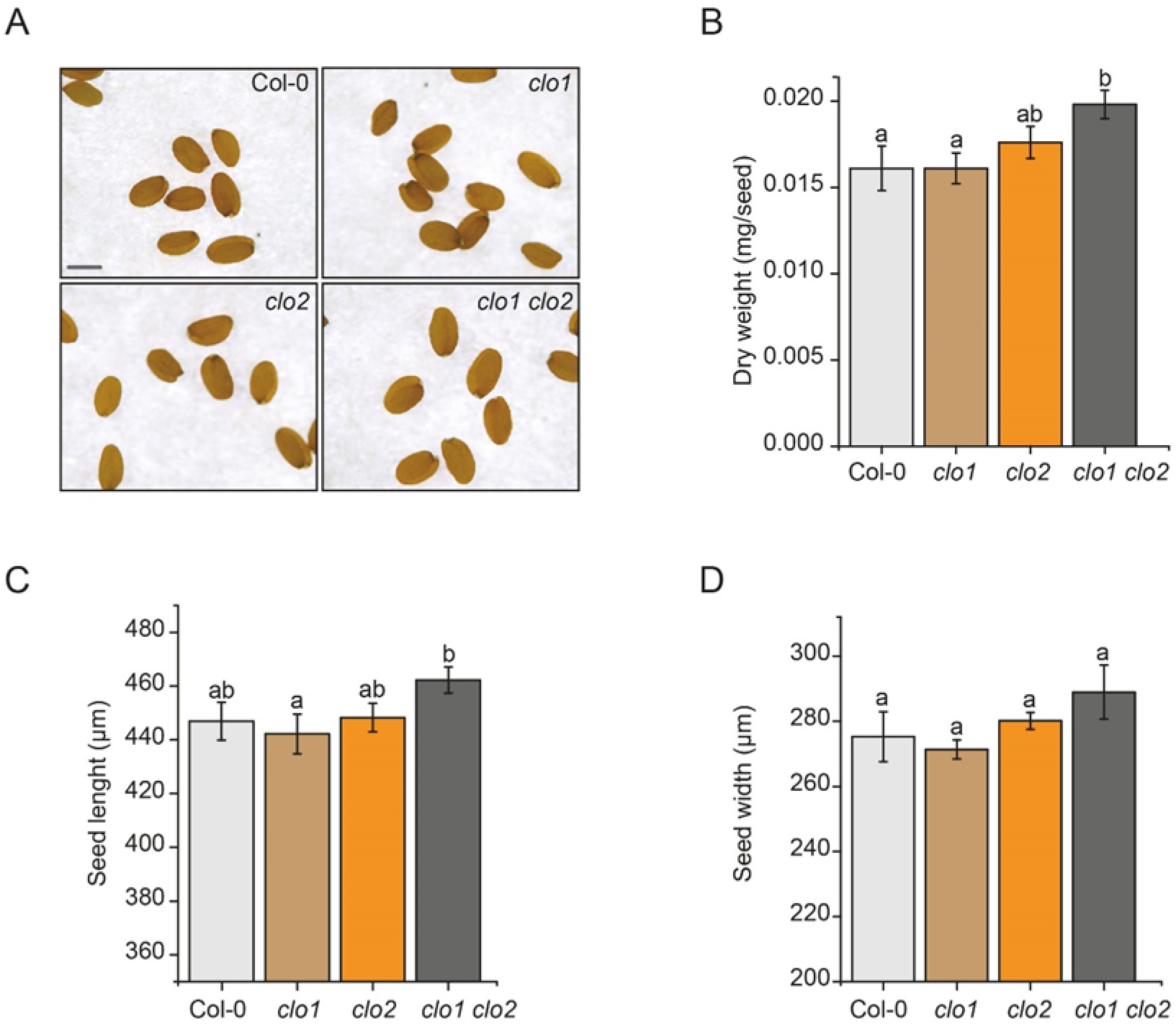
The effect of loss of function of CLO1 and CLO2 on Arabidopsis seed morphometric parameters. (A) Morphology of the Col-0 and *clo1*, *clo2* and *clo1 clo2* dry mature seeds, scale bar = 500 μm. (B) Dry weight of 300 mature seeds of Col-0, *clo1*, *clo2* and *clo1 clo2*. (C) Length of 150 mature seeds of Col-0, *clo1*, *clo2* and *clo1 clo2* (D) Width of 150 mature seeds of Col-0, *clo1*, *clo2* and *clo1 clo2*. Data are means ± SD from 9 biological replicates. Statistical analysis was performed by one-way ANOVA with Tukey‘s post hoc test. Different letters indicate significant differences with P < 0.05.

**Supplemental Figure S3.**
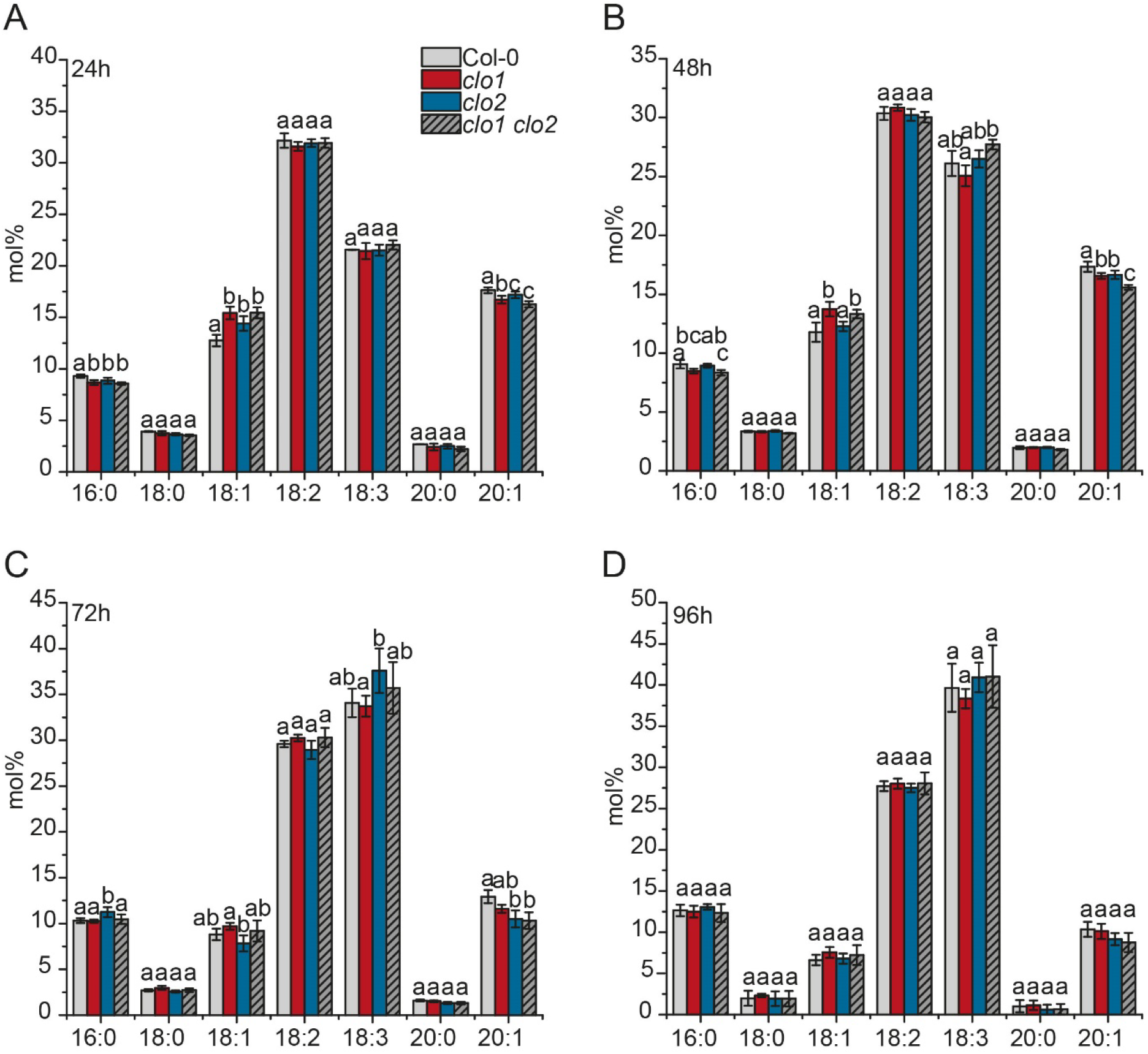
Fatty acid analysis in Col-0 and caleosin mutants during seed germination under long day conditions. (A-D) Changes in FAs (mol%) composition between Col-0, *clo1*, *clo2*, *clo1 clo2* after 24 h (A), 48 h (B), 72 h (C) and 96 h (D) of seed germination. Data are means ± SD from two independent experiments of six biological replicates (n = 6). Statistical analysis was performed by one-way ANOVA with Tukey‘s post hoc test. Different letters indicate significant differences with P < 0.05.

**Supplemental Figure S4.**
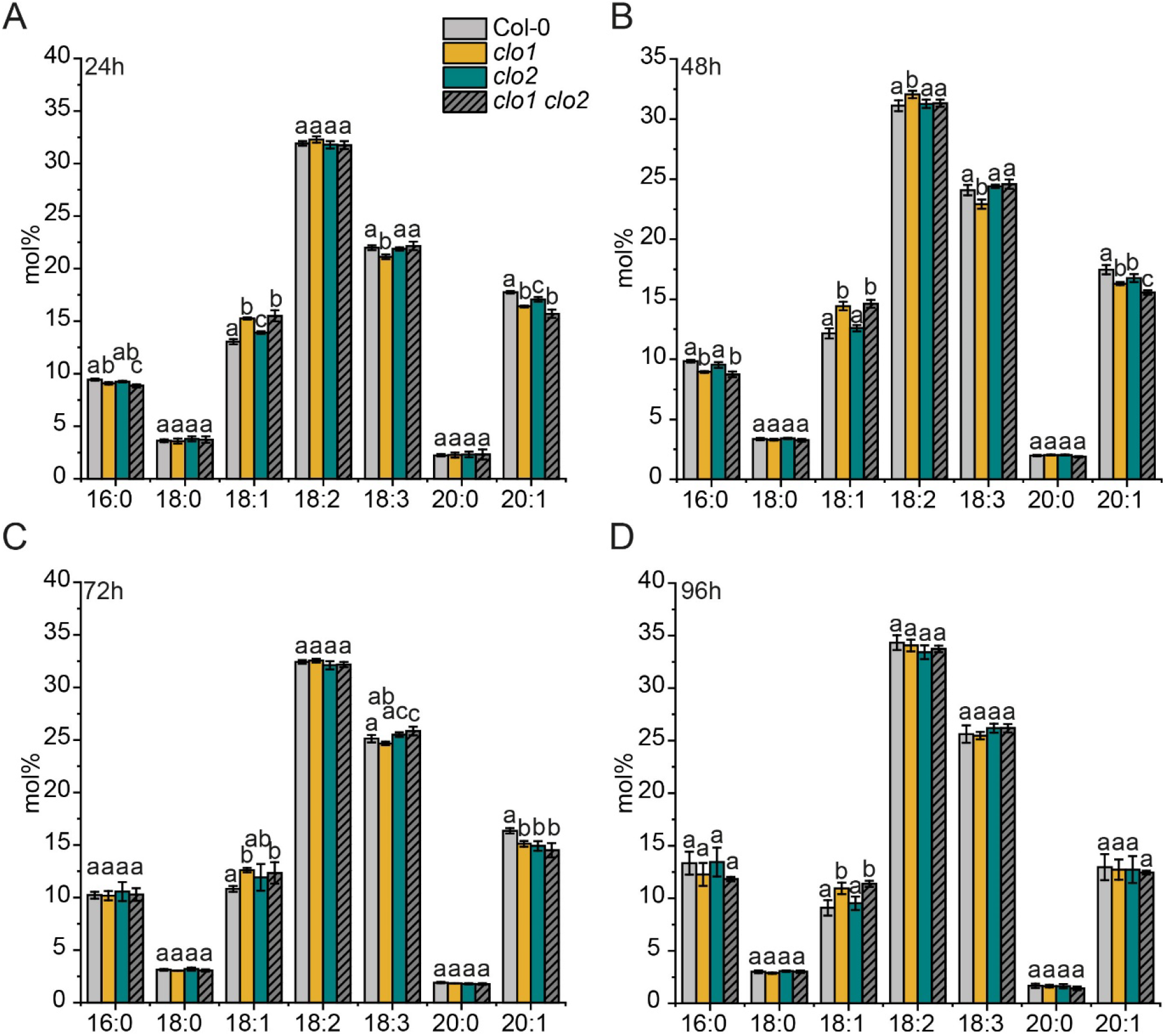
Fatty acid analysis of Col-0 and caleosin mutants during seed germination under continuous dark conditions. (A-D) Changes in FAs (mol%) composition between Col-0, clo1, clo2, clo1 clo2 after 24 h (A) 48 h (B), 72 h (C) and 96 h (D) of seed germination. Data are means ± SD from two independent experiments of six biological replicates (n = 6). Statistical analysis was performed by one-way ANOVA with Tukey‘s post hoc test. Different letters indicate significant differences with P < 0.05.

**Supplemental Figure S5.**
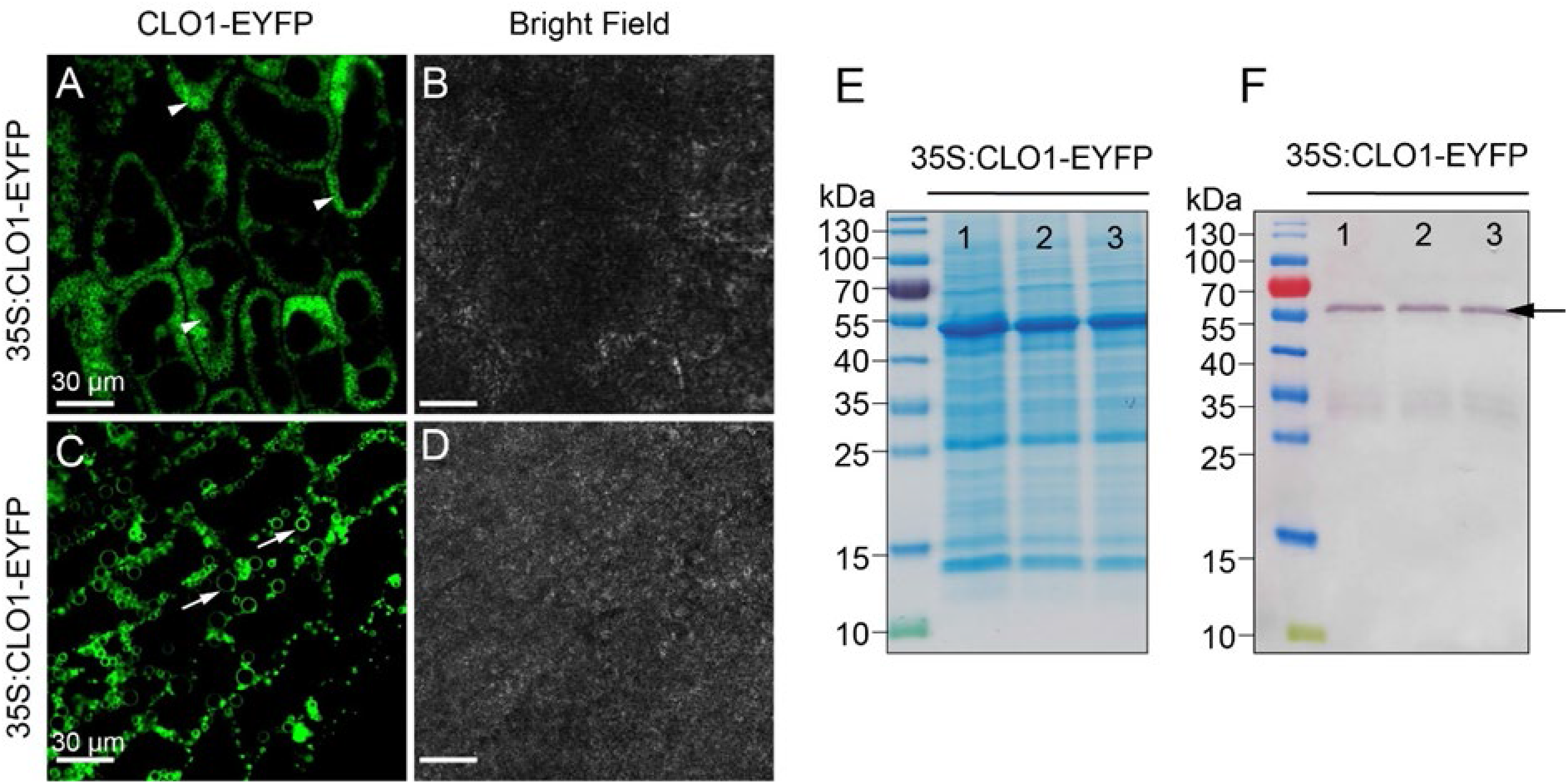
Expression of CLO1-EYFP in *A. thaliana* transgenic plants. (A-D) Representative CLSM images of *A. thaliana* seeds expressing CLO1-EYFP (green) under 35S promoter after 24 h (A-B) and 48 h (C-D) of seed germination. Arrowheads indicate CLO1-EYFP localized to cytoplasmic LDs. Arrows indicate the vacuolar pool of LDs. (E) Coomassie brilliant blue stained SDS-PAGE gel of total proteins from the 4-week old rosette leaves of three *A. thaliana* plants (lines 1, 2, 3) expressing CLO1-EYFP. (F) Detection of CLO1-EYFP by immunoblotting in 4-week old rosette leaves of three *A. thaliana* transgenic plants (lines 1, 2, 3). The arrow indicates CLO1-EYFP.

**Supplemental Figure S6.**
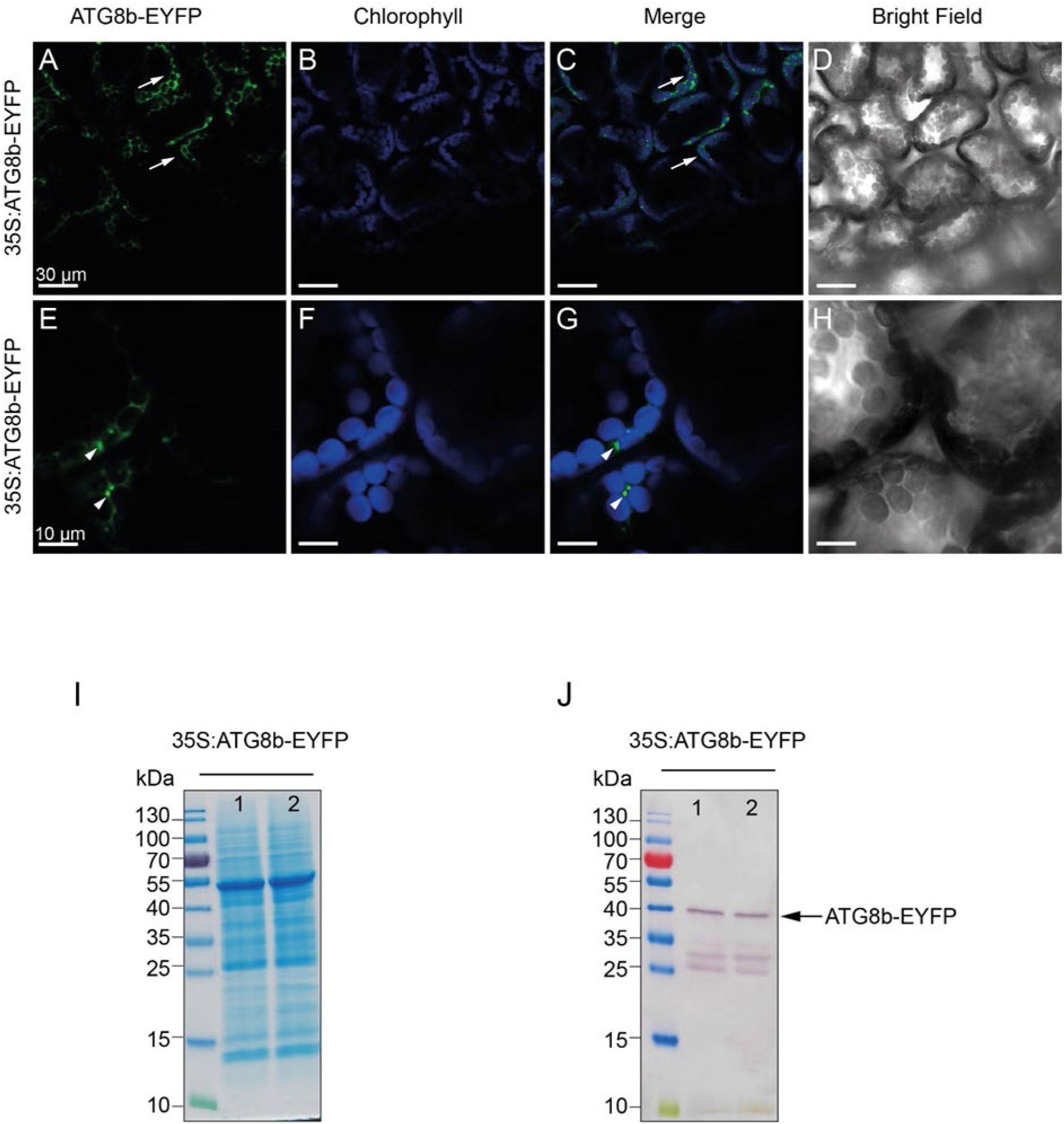
Expression of ATG8b-EYFP in *A. thaliana* transgenic plants. (A-H) Representative CLSM images of 4-week old rosette leaves of *A. thaliana* expressing ATG8b-EYFP (green) under 35S promoter. Arrows indicate cytoplasmic pool of ATG8b and arrowheads indicate ATG8b–labelled autophagic structures (I) SDS-PAGE gel of total proteins isolated from 4-week old rosette leaves and stained by Coomassie blue. (J) Immunoblotting detection of ATG8b-EYFP by using anti-GFP antibody. Line 1 and 2 correspond to two biological replicates.

**Supplemental Figure S7.**
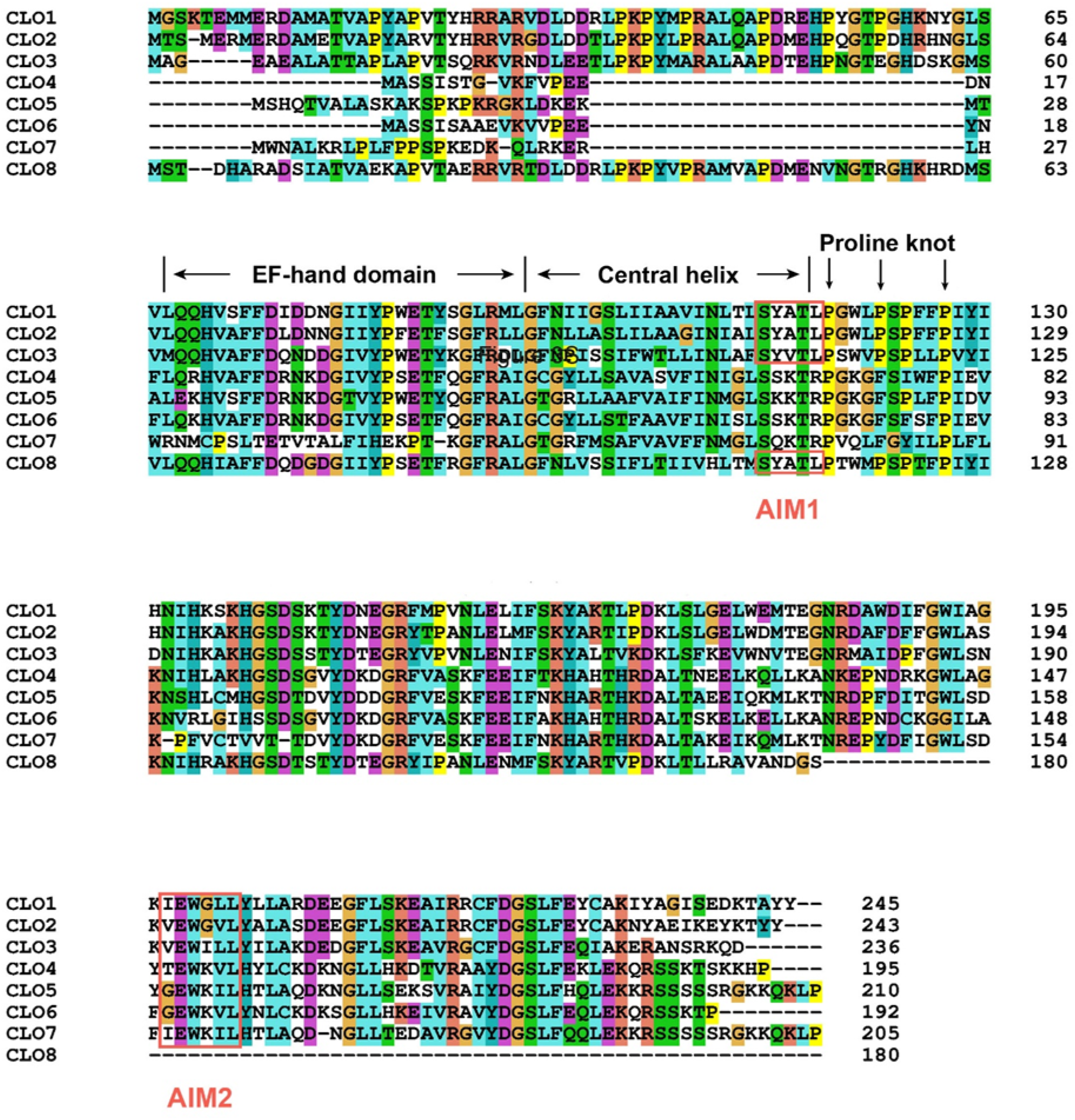
Multiple alignment of *A. thaliana* caleosins. The protein sequences of *A. thaliana* caleosins with the following accession numbers: CLO1 (O81270); CLO2 (Q9FLN9); CLO3 (O22788); CLO4 (Q9CAB7); CLO5 (B3H7A9); CLO6 (Q9CAB8); CLO7 (F4I4P8); CLO8 (A0A178V895) were obtained from UniProt database (www.uniprot.org) and aligned by ClustalW using MEGA-X software. The EF-hand motif, the central helix and the proline knot are shown above the sequences according to Purkrtova et al. (2007). Putative **A**TG8-**I**nteracting **M**otifs - AIMs (AIM1 and AIM2) are highlighted by pink frames.

**Supplemental Figure S8.**
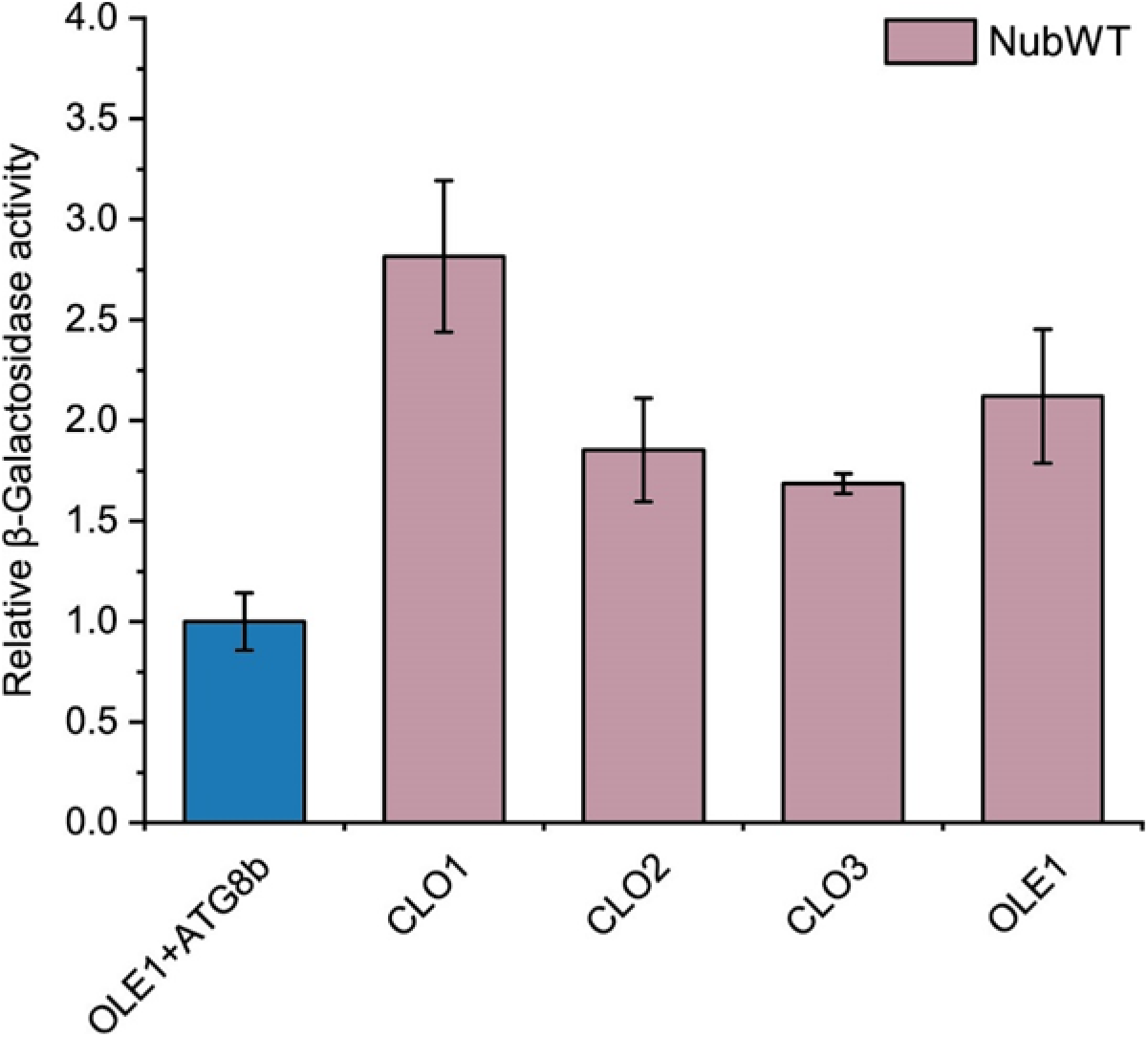
Interaction between caleosins and NubWT. Quantitative β-galactosidase activity assay for the Cub fusions of CLO1, CLO2, CLO3 or OLE1 with NubWT (positive control). Data are means ± SD from 4 independent yeast transformants. The β-galactosidase activity was normalized relative to the activity measured for the interaction between OLE1 and ATG8b (blue bar).

**Supplemental Figure S9.**
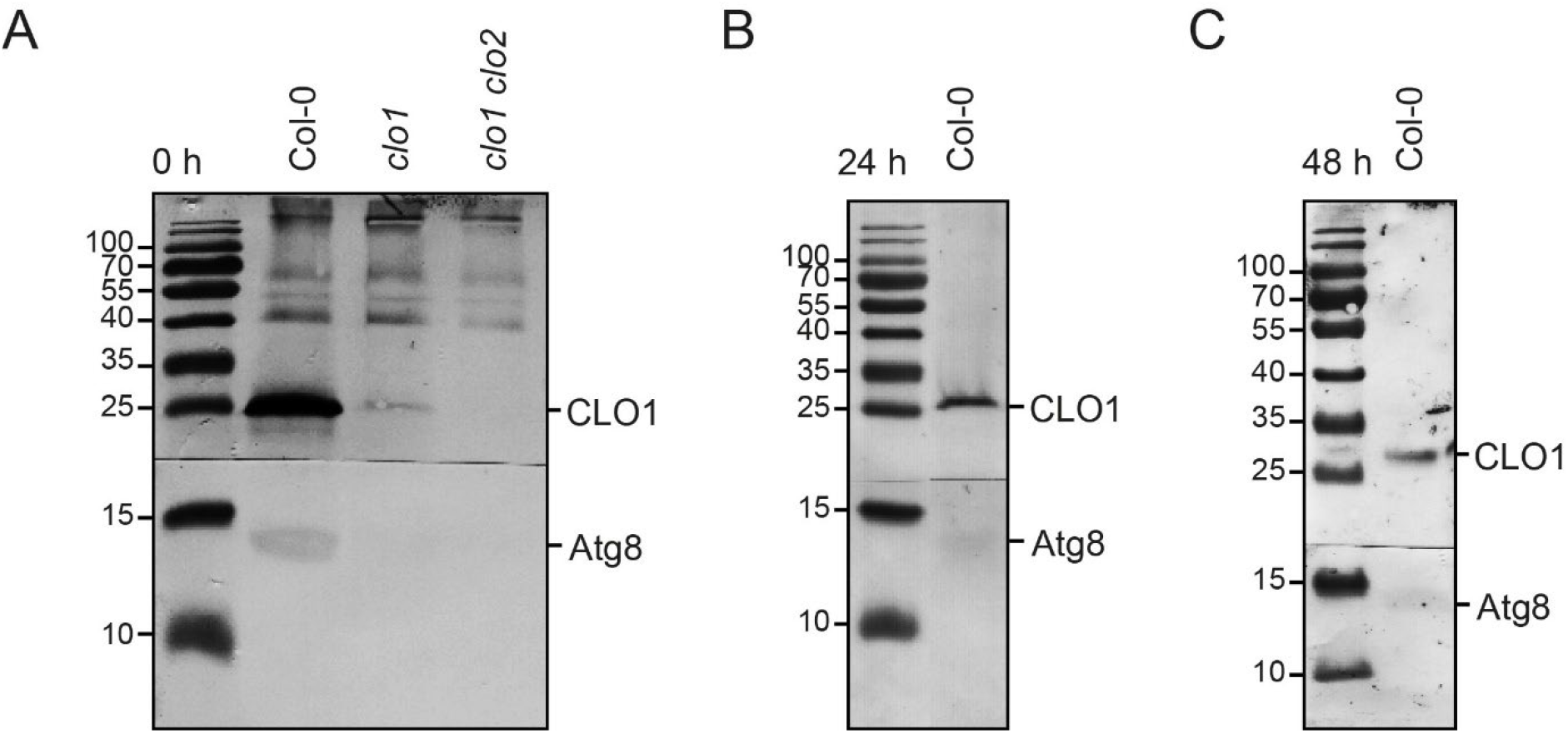
Co-immunoprecipitation assay showing the binding of CLO1 to ATG8. Proteins extracted from LD-fraction isolated from Col-0, *clo1*, *clo1 clo2* imbibed seeds (0 h) (A) and Col-0 seeds at 24 h (B) and at 48 h (C) of germination were immunoprecipitated using an anti-CLO1 antibody. The co-immunoprecipitated proteins were detected using an anti-CLO1 antibody (the upper part of the blot) and an anti-Atg8 antibody (the lower part of the blot).

**Supplemental Data Set 1. Raw data from lipid analysis and quantitative β-galactosidase assays.**

**Supplemental Data Set 2. ANOVA and Student’s t test results**

