## Supplemental DataSet 1 and 2 for "Caleosin 1 contributes to seed lipid droplet degradation by interaction with autophagy-related protein ATG8": Suplemental Dataset 2_Miklaszwewska et al., 2022.docx

| Anova Table | Sum of squares (SS) | Degrees of  freedom ν | Mean square  MS | F statistic | p-value |
| --- | --- | --- | --- | --- | --- |
| Treatment | 1.1689 | 3 | 0.3896 | 5.4604 | 0.0065 |
| Error | 1.4245 | 20 | 0.0712 |  |  |
| Total | 2.5934 | 23 |  | |  |

Supplemental File 2. ANOVA Tables

**Figure 1A**

a) 16:0
b) 18:0

| Anova Table | Sum of squares (SS) | Degrees of  freedom ν | Mean square  MS | F statistic | p-value |
| --- | --- | --- | --- | --- | --- |
| Treatment | 0.1840 | 3 | 0.0613 | 3.8737 | 0.0247 |
| Error | 0.3167 | 20 | 0.0158 |  |  |
| Total | 0.5007 | 23 |  | |  |

c) 18:1

| Anova Table | Sum of squares (SS) | Degrees of  freedom ν | Mean square  MS | F statistic | p-value |
| --- | --- | --- | --- | --- | --- |
| Treatment | 24.5792 | 3 | 8.1931 | 24.8262 | 6.0082e-07 |
| Error | 6.6003 | 20 | 0.3300 |  |  |
| Total | 31.1795 | 23 |  | |  |

d) 18:2

| Anova Table | Sum of squares (SS) | Degrees of  freedom ν | Mean square  MS | F statistic | p-value |
| --- | --- | --- | --- | --- | --- |
| Treatment | 1.3558 | 3 | 0.4519 | 1.2951 | 0.3035 |
| Error | 6.9787 | 20 | 0.3489 |  |  |
| Total | 8.3345 | 23 |  | |  |

e) 18:3

| Anova Table | Sum of squares (SS) | Degrees of  freedom ν | Mean square  MS | F statistic | p-value |
| --- | --- | --- | --- | --- | --- |
| Treatment | 3.8550 | 3 | 1.2850 | 6.0520 | 0.0042 |
| Error | 4.2465 | 20 | 0.2123 |  |  |
| Total | 8.1015 | 23 |  | |  |

f) 20:0

| Anova Table | Sum of squares (SS) | Degrees of  freedom ν | Mean square  MS | F statistic | p-value |
| --- | --- | --- | --- | --- | --- |
| Treatment | 0.1111 | 3 | 0.0370 | 5.7149 | 0.0054 |
| Error | 0.1296 | 20 | 0.0065 |  |  |
| Total | 0.2407 | 23 |  | |  |

g) 20:1

| Anova Table | Sum of squares (SS) | Degrees of  freedom ν | Mean square  MS | F statistic | p-value |
| --- | --- | --- | --- | --- | --- |
| Treatment | 13.6347 | 3 | 4.5449 | 29.6994 | 1.4479e-07 |
| Error | 3.0606 | 20 | 0.1530 |  |  |
| Total | 16.6953 | 23 |  | |  |

**Figure 1B**

a) 24h

| Anova Table | Sum of squares (SS) | Degrees of  freedom ν | Mean square  MS | F statistic | p-value |
| --- | --- | --- | --- | --- | --- |
| Treatment | 38.3669 | 3 | 12.7890 | 1.3649 | 0.2822 |
| Error | 187.4005 | 20 | 9.3700 |  |  |
| Total | 225.7673 | 23 |  | |  |

b) 48h

| Anova Table | Sum of squares (SS) | Degrees of  freedom ν | Mean square  MS | F statistic | p-value |
| --- | --- | --- | --- | --- | --- |
| Treatment | 76.7401 | 3 | 25.5800 | 2.0509 | 0.1408 |
| Error | 236.9813 | 19 | 12.4727 |  |  |
| Total | 313.7213 | 22 |  | |  |

c) 72h

| Anova Table | Sum of squares (SS) | Degrees of  freedom ν | Mean square  MS | F statistic | p-value |
| --- | --- | --- | --- | --- | --- |
| Treatment | 497.1991 | 3 | 165.7330 | 6.7225 | 0.0026 |
| Error | 493.0676 | 20 | 24.6534 |  |  |
| Total | 990.2667 | 23 |  | |  |

d) 96h

| Anova Table | Sum of squares (SS) | Degrees of  freedom ν | Mean square  MS | F statistic | p-value |
| --- | --- | --- | --- | --- | --- |
| Treatment | 332.1191 | 3 | 110.7064 | 4.7138 | 0.0353 |
| Error | 187.8860 | 19 | 23.4857 |  |  |
| Total | 520.0050 | 22 |  | |  |

**Figure 1C**

a) 24h

| Anova Table | Sum of squares (SS) | Degrees of  freedom ν | Mean square  MS | F statistic | p-value |
| --- | --- | --- | --- | --- | --- |
| Treatment | 15.225 | 3 | 5.0750 | 0.7917 | 0.5128 |
| Error | 128.2106 | 20 | 6.4105 |  |  |
| Total | 143.4357 | 23 |  | |  |

b) 48h

| Anova Table | Sum of squares (SS) | Degrees of  freedom ν | Mean square  MS | F statistic | p-value |
| --- | --- | --- | --- | --- | --- |
| Treatment | 24.3929 | 3 | 8.1310 | 0.7881 | 0.5146 |
| Error | 206.3386 | 20 | 10.3169 |  |  |
| Total | 230.7315 | 23 |  | |  |

c) 72h

| Anova Table | Sum of squares (SS) | Degrees of  freedom ν | Mean square  MS | F statistic | p-value |
| --- | --- | --- | --- | --- | --- |
| Treatment | 117.6361 | 3 | 39.2120 | 2.6124 | 0.076 |
| Error | 300.2029 | 20 | 15.0101 |  |  |
| Total | 417.8390 | 23 |  | |  |

d) 96h

| Anova Table | Sum of squares (SS) | Degrees of  freedom ν | Mean square  MS | F statistic | p-value |
| --- | --- | --- | --- | --- | --- |
| Treatment | 356.8523 | 3 | 118.9508 | 4.0495 | 0.0220 |
| Error | 558.1089 | 19 | 29.3742 |  |  |
| Total | 914.9612 | 22 |  | |  |

**Supplemental Figure 3S**

**Figure 3SA**

a) 16:0

| Anova Table | Sum of squares (SS) | Degrees of  freedom ν | Mean square  MS | F statistic | p-value |
| --- | --- | --- | --- | --- | --- |
| Treatment | 1.8415 | 3 | 0.6138 | 9.9965 | 0.0003 |
| Error | 1.2281 | 20 | 0.0614 |  |  |
| Total | 3.0697 | 23 |  | |  |

b) 18:0

| Anova Table | Sum of squares (SS) | Degrees of  freedom ν | Mean square  MS | F statistic | p-value |
| --- | --- | --- | --- | --- | --- |
| Treatment | 0.4314 | 3 | 0.1438 | 0.8981 | 0.4594 |
| Error | 3.2025 | 20 | 0.1601 |  |  |
| Total | 3.6340 | 23 |  | |  |

c) 18:1

| Anova Table | Sum of squares (SS) | Degrees of  freedom ν | Mean square  MS | F statistic | p-value |
| --- | --- | --- | --- | --- | --- |
| Treatment | 28.6311 | 3 | 9.5437 | 23.1982 | 1.0149e-06 |
| Error | 8.2280 | 20 | 0.4114 |  |  |
| Total | 36.8591 | 23 |  | |  |

d) 18:2

| Anova Table | Sum of squares (SS) | Degrees of  freedom ν | Mean square  MS | F statistic | p-value |
| --- | --- | --- | --- | --- | --- |
| Treatment | 0.9612 | 3 | 0.3204 | 0.5914 | 0.6278 |
| Error | 10.8361 | 20 | 0.5418 |  |  |
| Total | 11.7973 | 23 |  | |  |

e) 18:3

| Anova Table | Sum of squares (SS) | Degrees of  freedom ν | Mean square  MS | F statistic | p-value |
| --- | --- | --- | --- | --- | --- |
| Treatment | 1.3124 | 3 | 0.4375 | 0.9519 | 0.4345 |
| Error | 9.1916 | 20 | 0.4596 |  |  |
| Total | 10.5040 | 23 |  | |  |

f) 20:0

| Anova Table | Sum of squares (SS) | Degrees of  freedom ν | Mean square  MS | F statistic | p-value |
| --- | --- | --- | --- | --- | --- |
| Treatment | 0.6602 | 3 | 0.2201 | 1.0995 | 0.3725 |
| Error | 4.0034 | 20 | 0.2002 |  |  |
| Total | 4.6636 | 23 |  | |  |

g) 20:1

| Anova Table | Sum of squares (SS) | Degrees of  freedom ν | Mean square  MS | F statistic | p-value |
| --- | --- | --- | --- | --- | --- |
| Treatment | 6.2431 | 3 | 2.0810 | 11.7780 | 0.0001 |
| Error | 3.5337 | 20 | 0.1767 |  |  |
| Total | 9.7768 | 23 |  | |  |

**Figure 3SB**

a) 16:0

| Anova Table | Sum of squares (SS) | Degrees of  freedom ν | Mean square  MS | F statistic | p-value |
| --- | --- | --- | --- | --- | --- |
| Treatment | 2.2274 | 3 | 0.7425 | 9.6047 | 0.0005 |
| Error | 1.4688 | 19 | 0.0773 |  |  |
| Total | 3.6962 | 22 |  | |  |

b) 18:0

| Anova Table | Sum of squares (SS) | Degrees of  freedom ν | Mean square  MS | F statistic | p-value |
| --- | --- | --- | --- | --- | --- |
| Treatment | 0.1603 | 3 | 0.0534 | 11.2548 | 0.0002 |
| Error | 0.0902 | 19 | 0.0047 |  |  |
| Total | 0.2505 | 22 |  | |  |

c) 18:1

| Anova Table | Sum of squares (SS) | Degrees of  freedom ν | Mean square  MS | F statistic | p-value |
| --- | --- | --- | --- | --- | --- |
| Treatment | 13.9820 | 3 | 4.6607 | 11.4987 | 0.0002 |
| Error | 7.7011 | 19 | 0.4053 |  |  |
| Total | 21.6831 | 22 |  | |  |

d) 18:2

| Anova Table | Sum of squares (SS) | Degrees of  freedom ν | Mean square  MS | F statistic | p-value |
| --- | --- | --- | --- | --- | --- |
| Treatment | 1.9632 | 3 | 0.6544 | 2.5131 | 0.0893 |
| Error | 4.9476 | 19 | 0.2604 |  |  |
| Total | 6.9108 | 22 |  | |  |

e) 18:3

| Anova Table | Sum of squares (SS) | Degrees of  freedom ν | Mean square  MS | F statistic | p-value |
| --- | --- | --- | --- | --- | --- |
| Treatment | 20.1029 | 3 | 6.7010 | 8.4329 | 0.0009 |
| Error | 15.0978 | 19 | 0.7946 |  |  |
| Total | 35.2007 | 22 |  | |  |

f) 20:0

| Anova Table | Sum of squares (SS) | Degrees of  freedom ν | Mean square  MS | F statistic | p-value |
| --- | --- | --- | --- | --- | --- |
| Treatment | 0.1431 | 3 | 0.0477 | 6.1643 | 0.0042 |
| Error | 0.1471 | 19 | 0.0077 |  |  |
| Total | 0.2902 | 22 |  | |  |

g) 20:1

| Anova Table | Sum of squares (SS) | Degrees of  freedom ν | Mean square  MS | F statistic | p-value |
| --- | --- | --- | --- | --- | --- |
| Treatment | 9.5166 | 3 | 3.1722 | 24.8584 | 8.6280e-07 |
| Error | 2.4246 | 19 | 0.1276 |  |  |
| Total | 11.9412 | 22 |  | |  |

**Figure 3SC**

a) 16:0

| Anova Table | Sum of squares (SS) | Degrees of  freedom ν | Mean square  MS | F statistic | p-value |
| --- | --- | --- | --- | --- | --- |
| Treatment | 3.7520 | 3 | 1.2507 | 6.3322 | 0.0034 |
| Error | 3.9502 | 20 | 0.1975 |  |  |
| Total | 7.7022 | 23 |  | |  |

b) 18:0

| Anova Table | Sum of squares (SS) | Degrees of  freedom ν | Mean square  MS | F statistic | p-value |
| --- | --- | --- | --- | --- | --- |
| Treatment | 0.4677 | 3 | 0.1559 | 4.6801 | 0.0124 |
| Error | 0.6662 | 20 | 0.0333 |  |  |
| Total | 1.1340 | 23 |  | |  |

c) 18:1

| Anova Table | Sum of squares (SS) | Degrees of  freedom ν | Mean square  MS | F statistic | p-value |
| --- | --- | --- | --- | --- | --- |
| Treatment | 11.4136 | 3 | 3.8045 | 4.7234 | 0.0119 |
| Error | 16.1093 | 20 | 0.8055 |  |  |
| Total | 27.5229 | 23 |  | |  |

d) 18:2

| Anova Table | Sum of squares (SS) | Degrees of  freedom ν | Mean square  MS | F statistic | p-value |
| --- | --- | --- | --- | --- | --- |
| Treatment | 7.3508 | 3 | 2.4503 | 3.4343 | 0.0366 |
| Error | 14.2696 | 20 | 0.7135 |  |  |
| Total | 21.6204 | 23 |  | |  |

e) 18:3

| Anova Table | Sum of squares (SS) | Degrees of  freedom ν | Mean square  MS | F statistic | p-value |
| --- | --- | --- | --- | --- | --- |
| Treatment | 56.6024 | 3 | 18.8675 | 3.5796 | 0.0321 |
| Error | 105.4166 | 20 | 5.2708 |  |  |
| Total | 162.0190 | 23 |  | |  |

f) 20:0

| Anova Table | Sum of squares (SS) | Degrees of  freedom ν | Mean square  MS | F statistic | p-value |
| --- | --- | --- | --- | --- | --- |
| Treatment | 0.3650 | 3 | 0.1217 | 6.6957 | 0.0026 |
| Error | 0.3634 | 20 | 0.0182 |  |  |
| Total | 0.7283 | 23 |  | |  |

g) 20:1

| Anova Table | Sum of squares (SS) | Degrees of  freedom ν | Mean square  MS | F statistic | p-value |
| --- | --- | --- | --- | --- | --- |
| Treatment | 26.1942 | 3 | 8.7314 | 12.2490 | 9.0181e-05 |
| Error | 14.2565 | 20 | 0.7128 |  |  |
| Total | 40.4507 | 23 |  | |  |

**Figure 3SD**

a) 16:0

| Anova Table | Sum of squares (SS) | Degrees of  freedom ν | Mean square  MS | F statistic | p-value |
| --- | --- | --- | --- | --- | --- |
| Treatment | 1.7666 | 3 | 0.5889 | 0.8358 | 0.4908 |
| Error | 13.3874 | 19 | 0.7046 |  |  |
| Total | 15.1541 | 22 |  | |  |

b) 18:0

| Anova Table | Sum of squares (SS) | Degrees of  freedom ν | Mean square  MS | F statistic | p-value |
| --- | --- | --- | --- | --- | --- |
| Treatment | 0.5272 | 3 | 0.1757 | 0.2185 | 0.8823 |
| Error | 15.2767 | 19 | 0.8040 |  |  |
| Total | 15.8039 | 22 |  | |  |

c) 18:1

| Anova Table | Sum of squares (SS) | Degrees of  freedom ν | Mean square  MS | F statistic | p-value |
| --- | --- | --- | --- | --- | --- |
| Treatment | 2.8966 | 3 | 0.9655 | 1.1972 | 0.3376 |
| Error | 15.3229 | 19 | 0.8065 |  |  |
| Total | 18.2195 | 22 |  | |  |

d) 18:2

| Anova Table | Sum of squares (SS) | Degrees of  freedom ν | Mean square  MS | F statistic | p-value |
| --- | --- | --- | --- | --- | --- |
| Treatment | 1.2472 | 3 | 0.4157 | 0.4759 | 0.7028 |
| Error | 16.6001 | 19 | 0.8737 |  |  |
| Total | 17.8473 | 22 |  | |  |

e) 18:3

| Anova Table | Sum of squares (SS) | Degrees of  freedom ν | Mean square  MS | F statistic | p-value |
| --- | --- | --- | --- | --- | --- |
| Treatment | 25.8427 | 3 | 8.6142 | 1.0040 | 0.4126 |
| Error | 163.0155 | 19 | 8.5798 |  |  |
| Total | 188.8582 | 22 |  | |  |

f) 20:0

| Anova Table | Sum of squares (SS) | Degrees of  freedom ν | Mean square  MS | F statistic | p-value |
| --- | --- | --- | --- | --- | --- |
| Treatment | 1.1803 | 3 | 0.3934 | 0.7693 | 0.5254 |
| Error | 9.7174 | 19 | 0.5114 |  |  |
| Total | 10.8977 | 22 |  | |  |

g) 20:1

| Anova Table | Sum of squares (SS) | Degrees of  freedom ν | Mean square  MS | F statistic | p-value |
| --- | --- | --- | --- | --- | --- |
| Treatment | 9.9915 | 3 | 3.3305 | 3.0657 | 0.0529 |
| Error | 20.6412 | 19 | 1.0864 |  |  |
| Total | 30.6327 | 22 |  | |  |

**Supplemental Figure 4S**

**Figure 4SA**

a) 16:0

| Anova Table | Sum of squares (SS) | Degrees of  freedom ν | Mean square  MS | F statistic | p-value |
| --- | --- | --- | --- | --- | --- |
| Treatment | 1.1596 | 3 | 0.3865 | 25.7042 | 4.5786e-07 |
| Error | 0.3008 | 20 | 0.0150 |  |  |
| Total | 1.4604 | 23 |  | |  |

b) 18:0

| Anova Table | Sum of squares (SS) | Degrees of  freedom ν | Mean square  MS | F statistic | p-value |
| --- | --- | --- | --- | --- | --- |
| Treatment | 0.1738 | 3 | 0.0579 | 0.8642 | 0.4759 |
| Error | 1.3408 | 20 | 0.0670 |  |  |
| Total | 1.5146 | 23 |  | |  |

c) 18:1

| Anova Table | Sum of squares (SS) | Degrees of  freedom ν | Mean square  MS | F statistic | p-value |
| --- | --- | --- | --- | --- | --- |
| Treatment | 23.9957 | 3 | 7.9986 | 71.8572 | 6.9158e-11 |
| Error | 2.2262 | 20 | 0.1113 |  |  |
| Total | 26.2219 | 23 |  | |  |

d) 18:2

| Anova Table | Sum of squares (SS) | Degrees of  freedom ν | Mean square  MS | F statistic | p-value |
| --- | --- | --- | --- | --- | --- |
| Treatment | 1.0821 | 3 | 0.3607 | 2.8866 | 0.0611 |
| Error | 2.4992 | 20 | 0.1250 |  |  |
| Total | 3.5813 | 23 |  | |  |

e) 18:3

| Anova Table | Sum of squares (SS) | Degrees of  freedom ν | Mean square  MS | F statistic | p-value |
| --- | --- | --- | --- | --- | --- |
| Treatment | 3.7439 | 3 | 1.2480 | 13.7756 | 4.2225e-05 |
| Error | 1.8118 | 20 | 0.0906 |  |  |
| Total | 5.5557 | 23 |  | |  |

f) 20:0

| Anova Table | Sum of squares (SS) | Degrees of  freedom ν | Mean square  MS | F statistic | p-value |
| --- | --- | --- | --- | --- | --- |
| Treatment | 0.0423 | 3 | 0.0141 | 0.1275 | 0.9427 |
| Error | 2.2090 | 20 | 0.1105 |  |  |
| Total | 2.2513 | 23 |  | |  |

g) 20:1

| Anova Table | Sum of squares (SS) | Degrees of  freedom ν | Mean square  MS | F statistic | p-value |
| --- | --- | --- | --- | --- | --- |
| Treatment | 14.2145 | 3 | 4.7382 | 69.6566 | 9.1784e-11 |
| Error | 1.3604 | 20 | 0.0680 |  |  |
| Total | 15.5749 | 23 |  | |  |

**Figure 4SB**

a) 16:0

| Anova Table | Sum of squares (SS) | Degrees of  freedom ν | Mean square  MS | F statistic | p-value |
| --- | --- | --- | --- | --- | --- |
| Treatment | 4.7272 | 3 | 4.7382 | 69.6566 | 9.1784e-11 |
| Error | 0.7759 | 20 | 0.0680 |  |  |
| Total | 5.5032 | 23 |  | |  |

b) 18:0

| Anova Table | Sum of squares (SS) | Degrees of  freedom ν | Mean square  MS | F statistic | p-value |
| --- | --- | --- | --- | --- | --- |
| Treatment | 0.0755 | 3 | 0.0252 | 2.7292 | 0.0711 |
| Error | 0.1845 | 20 | 0.0092 |  |  |
| Total | 0.2600 | 23 |  | |  |

c) 18:1

| Anova Table | Sum of squares (SS) | Degrees of  freedom ν | Mean square  MS | F statistic | p-value |
| --- | --- | --- | --- | --- | --- |
| Treatment | 28.3943 | 3 | 9.4648 | 66.3134 | 1.4336e-10 |
| Error | 2.8546 | 20 | 0.1427 |  |  |
| Total | 31.2489 | 23 |  | |  |

d) 18:2

| Anova Table | Sum of squares (SS) | Degrees of  freedom ν | Mean square  MS | F statistic | p-value |
| --- | --- | --- | --- | --- | --- |
| Treatment | 3.0880 | 3 | 1.0293 | 6.6912 | 0.0026 |
| Error | 3.0766 | 20 | 0.1538 |  |  |
| Total | 6.1646 | 23 |  | |  |

e) 18:3

| Anova Table | Sum of squares (SS) | Degrees of  freedom ν | Mean square  MS | F statistic | p-value |
| --- | --- | --- | --- | --- | --- |
| Treatment | 10.1191 | 3 | 3.3730 | 22.7823 | 1.1656e-06 |
| Error | 2.9611 | 20 | 0.1481 |  |  |
| Total | 13.0802 | 23 |  | |  |

f) 20:0

| Anova Table | Sum of squares (SS) | Degrees of  freedom ν | Mean square  MS | F statistic | p-value |
| --- | --- | --- | --- | --- | --- |
| Treatment | 0.0819 | 3 | 0.0273 | 8.1465 | 0.0010 |
| Error | 0.0671 | 20 | 0.0034 |  |  |
| Total | 0.1490 | 23 |  | |  |

g) 20:1

| Anova Table | Sum of squares (SS) | Degrees of  freedom ν | Mean square  MS | F statistic | p-value |
| --- | --- | --- | --- | --- | --- |
| Treatment | 11.3892 | 3 | 3.7964 | 40.0849 | 1.2000e-08 |
| Error | 1.8942 | 20 | 0.0947 |  |  |
| Total | 13.2834 | 23 |  | |  |

**Figure 4SC**

a) 16:0

| Anova Table | Sum of squares (SS) | Degrees of  freedom ν | Mean square  MS | F statistic | p-value |
| --- | --- | --- | --- | --- | --- |
| Treatment | 0.5516 | 3 | 0.1839 | 0.4162 | 0.7433 |
| Error | 8.8359 | 20 | 0.4418 |  |  |
| Total | 9.3875 | 23 |  | |  |

b) 18:0

| Anova Table | Sum of squares (SS) | Degrees of  freedom ν | Mean square  MS | F statistic | p-value |
| --- | --- | --- | --- | --- | --- |
| Treatment | 0.0872 | 3 | 0.0291 | 2.5138 | 0.0877 |
| Error | 0.2313 | 20 | 0.0116 |  |  |
| Total | 0.3185 | 23 |  | |  |

c) 18:1

| Anova Table | Sum of squares (SS) | Degrees of  freedom ν | Mean square  MS | F statistic | p-value |
| --- | --- | --- | --- | --- | --- |
| Treatment | 11.0432 | 3 | 3.6811 | 4.3982 | 0.0157 |
| Error | 16.7388 | 20 | 0.8369 |  |  |
| Total | 27.7820 | 23 |  | |  |

d) 18:2

| Anova Table | Sum of squares (SS) | Degrees of  freedom ν | Mean square  MS | F statistic | p-value |
| --- | --- | --- | --- | --- | --- |
| Treatment | 0.7926 | 3 | 0.2642 | 3.2409 | 0.0438 |
| Error | 1.6303 | 20 | 0.0815 |  |  |
| Total | 2.4229 | 23 |  | |  |

e) 18:3

| Anova Table | Sum of squares (SS) | Degrees of  freedom ν | Mean square  MS | F statistic | p-value |
| --- | --- | --- | --- | --- | --- |
| Treatment | 4.6415 | 3 | 1.5472 | 14.3918 | 3.1570e-05 |
| Error | 2.1501 | 20 | 0.1075 |  |  |
| Total | 6.7916 | 23 |  | |  |

f) 20:0

| Anova Table | Sum of squares (SS) | Degrees of  freedom ν | Mean square  MS | F statistic | p-value |
| --- | --- | --- | --- | --- | --- |
| Treatment | 0.0735 | 3 | 0.0245 | 5.7321 | 0.0053 |
| Error | 0.0855 | 20 | 0.0043 |  |  |
| Total | 0.1591 | 23 |  | |  |

g) 20:1

| Anova Table | Sum of squares (SS) | Degrees of  freedom ν | Mean square  MS | F statistic | p-value |
| --- | --- | --- | --- | --- | --- |
| Treatment | 11.4300 | 3 | 3.8100 | 15.5160 | 1.8959e-05 |
| Error | 4.9111 | 20 | 0.2456 |  |  |
| Total | 16.3411 | 23 |  | |  |

**Figure 4SD**

a) 16:0

| Anova Table | Sum of squares (SS) | Degrees of  freedom ν | Mean square  MS | F statistic | p-value |
| --- | --- | --- | --- | --- | --- |
| Treatment | 10.7215 | 3 | 3.5738 | 2.6201 | 0.0806 |
| Error | 25.9161 | 19 | 1.3640 |  |  |
| Total | 36.6375 | 22 |  | |  |

b) 18:0

| Anova Table | Sum of squares (SS) | Degrees of  freedom ν | Mean square  MS | F statistic | p-value |
| --- | --- | --- | --- | --- | --- |
| Treatment | 0.1142 | 3 | 0.0381 | 2.7205 | 0.0732 |
| Error | 0.2657 | 19 | 0.0140 |  |  |
| Total | 0.3799 | 22 |  | |  |

c) 18:1

| Anova Table | Sum of squares (SS) | Degrees of  freedom ν | Mean square  MS | F statistic | p-value |
| --- | --- | --- | --- | --- | --- |
| Treatment | 20.5240 | 3 | 6.8413 | 16.7676 | 1.4353e-05 |
| Error | 7.7522 | 19 | 0.4080 |  |  |
| Total | 28.2762 | 22 |  | |  |

d) 18:2

| Anova Table | Sum of squares (SS) | Degrees of  freedom ν | Mean square  MS | F statistic | p-value |
| --- | --- | --- | --- | --- | --- |
| Treatment | 2.8139 | 3 | 0.9380 | 2.2922 | 0.1108 |
| Error | 7.7749 | 19 | 0.4092 |  |  |
| Total | 10.5888 | 22 |  | |  |

e) 18:3

| Anova Table | Sum of squares (SS) | Degrees of  freedom ν | Mean square  MS | F statistic | p-value |
| --- | --- | --- | --- | --- | --- |
| Treatment | 2.3660 | 3 | 0.7887 | 2.1622 | 0.1260 |
| Error | 6.9304 | 19 | 0.3648 |  |  |
| Total | 9.2963 | 22 |  | |  |

f) 20:0

| Anova Table | Sum of squares (SS) | Degrees of  freedom ν | Mean square  MS | F statistic | p-value |
| --- | --- | --- | --- | --- | --- |
| Treatment | 0.1818 | 3 | 0.0606 | 1.4511 | 0.2594 |
| Error | 0.7936 | 19 | 0.0418 |  |  |
| Total | 0.9755 | 22 |  | |  |

g) 20:1

| Anova Table | Sum of squares (SS) | Degrees of  freedom ν | Mean square  MS | F statistic | p-value |
| --- | --- | --- | --- | --- | --- |
| Treatment | 0.7322 | 3 | 0.2441 | 0.1888 | 0.9027 |
| Error | 24.5584 | 19 | 1.2925 |  |  |
| Total | 25.2906 | 22 |  | |  |
